# Fine-scale tracking reveals visual field use for predator detection and escape in collective foraging of pigeon flocks

**DOI:** 10.1101/2024.02.05.578919

**Authors:** Mathilde Delacoux, Fumihiro Kano

**Affiliations:** Centre for the Advanced Study of Collective Behaviour, University of Konstanz, Germany; Max Planck Institute of Animal Behavior, Radolfzell, Germany; International Max Planck Research School for Quantitative Behavior, Ecology and Evolution, Radolfzell, Germany

**Keywords:** Fovea, predator detection, pigeon *(Columba livia)*, collective vigilance, social contagion

## Abstract

During collective vigilance, it is commonly assumed that individual animals compromise their feeding time to be vigilant against predators, benefiting the entire group. One notable issue with this assumption concerns the unclear nature of predator “detection”, particularly in terms of vision. It remains uncertain how a vigilant individual utilizes its high-acuity vision (such as the fovea) to detect a predator cue and subsequently guide individual and collective escape responses. Using fine-scale motion capture technologies, we tracked the head and body orientations of pigeons (hence reconstructed their visual fields and foveal projections) foraging in a flock during simulated predator attacks. Pigeons used their fovea to inspect predator cues. Earlier foveation on a predator cue was linked to preceding behaviors related to vigilance and feeding, such as head-up or down positions, head-scanning, and food-pecking. Moreover, earlier foveation predicted earlier evasion flights at both the individual and collective levels. However, we also found that relatively long delay between their foveation and escape responses in individuals obscured the relationship between these two responses. While our results largely support the existing assumptions about vigilance, they also underscore the importance of considering vision and addressing the disparity between detection and escape responses in future research.

## Introduction

In everyday natural tasks, such as locating food, avoiding predators, and interacting with conspecifics, animals constantly face challenges in deciding when and how to adjust their behaviors to increase their chances of survival (McFarland, 1977). One such behavior is the focusing of attention, or “looking” behavior, which has been relatively understudied in natural tasks due to the technical challenges involved in tracking an animal’s gaze during natural activities (but see Kane et al., 2015; Kano et al., 2018; Miñano et al., 2023; Yorzinski & Platt, 2014). However, this is relatively well studied within the context of vigilance, a scenario where an animal’s survival hinges on effective attentional allocation and where researchers can observe their scanning behavior even in field conditions (Cresswell, 1994; Evans et al., 2018; Inglis & Lazarus, 1981). Vigilance, especially during foraging, generally involves the compromise between staying alert for threats and searching for food, as well as maintaining a balance between individual and collective vigilance (Beauchamp, 2015).

### Common assumptions of vigilance research

Common assumptions about collective vigilance, first coined by Pulliam (Pulliam, 1973; Pulliam et al., 1982), posit that a vigilant individual, often identified by a “head-up” posture, is likely to have a higher probability of detecting an approaching predator compared to a feeding individual, typically in a “head-down” posture. Consequently, the vigilant individual can react and escape more swiftly, thereby reducing its predation risk (also see Beauchamp, 2015; Godin & Smith, 1988). In a group setting, non-vigilant members can gain benefits from the vigilant individual by following its lead when it exhibits escape behavior (Pulliam, 1973). This potential group advantage, known as “collective detection,” might explain that individuals in a larger group tend to be less vigilant (and hence spend more time in foraging), suggesting a critical advantage of grouping (Pulliam, 1973; Pulliam et al., 1982).

### Empirical challenges

While these assumptions underpin many influential theoretical models in behavioral ecology, they have not gone unchallenged, with numerous empirical studies highlighting potential inconsistencies. Firstly, the trade-off between vigilance and foraging may not be as pronounced as previously assumed. Even though several studies found a relationship between vigilance and escape latency (Devereux et al., 2006; Hilton et al., 1999; Lima & Bednekoff, 1999), various studies have demonstrated that these two behaviors are not necessarily mutually exclusive (Bednekoff & Lima, 2005; Cresswell et al., 2003; Devereux et al., 2006; Kaby & Lind, 2003; Lima & Bednekoff, 1999). In several species, especially those with a broad visual field and specific retinal structures such as the visual streaks, individuals can simultaneously engage in foraging activities while remaining vigilant (Fernández-Juricic, 2012), likely using peripheral vision to detect approaching threats (Bednekoff & Lima, 2005; Cresswell et al., 2003; Kaby & Lind, 2003; Lima & Bednekoff, 1999). Relatedly, although vigilance and foraging have been defined in many previous studies respectively as head-up and head-down due to the limitation associated with direct observation, several studies have pointed out that vigilance triggered by the appearance of a predator is not necessarily related to the head-up postures but to the pattern of head movements (Fernández-Juricic, 2012; Jones et al., 2007, 2009). Similarly, the patterns of head-up (number of head-up bouts, their length and regularity), rather than the proportion of time being head-up versus head-down, are sometimes better predictors of predator detection (Beauchamp, 2015; Beauchamp & Ruxton, 2016; Bednekoff & Lima, 2002; Cresswell et al., 2003; Hart & Lendrem, 1984).

Second, it is challenging to empirically differentiate the effect of collective detection from other group size-related effects, such as confusion, dilution, and edge effect. Although there is evidence that non-detectors can benefit from detectors’ escape (Davis, 1975; Lima, 1995b; Lima & Zollner, 1996), other studies found evidence for the risk dilution (Beauchamp & Ruxton, 2008) and the edge effect (Inglis & Lazarus, 1981) in their study systems. It appears that multiple factors influence vigilance across species, including group size, density, and spatial configuration, as well as social contagion (Elgar, 1989; Roberts, 1996).

Third, non-vigilant animals do not always respond to behavioral cues of other members of the group. The upright-freezing alert posture, often one of the first behavioral indications of predator detection, tends to be rather inconspicuous, and flockmates usually do not seem to respond to it (Fernández-Juricic et al., 2009; Lima, 1995b). Additionally, birds do not necessarily differentiate between predator-based escapes and non-threat departures from a flock mate, suggesting that simultaneous departures of multiple birds are required to trigger contagious flights (Cresswell et al., 2000; Lima, 1995b; Proctor et al., 2001). As a result, the extent to which group-living animals benefit from collective detection remains open to questions (Bednekoff & Lima, 1998).

Finally, the definition of “detection” is generally uncertain in studies. Previous studies typically used overt escape responses, such as flying, as a measure of detection (using escape as only measure: Kenward, 1978; Lima & Zollner, 1996; Quinn & Cresswell, 2005; or partially using escape as response: Cresswell et al., 2003; Lima, 1995a, 1995b; Tisdale & Fernández-Juricic, 2009; Whittingham et al., 2004). However, a bird likely identifies a potential threat before escaping (Barbosa & Castellanos, 2005; Fernández-Juricic, 2012; Lima & Dill, 1990). Thus, subtler behavioral responses like freezing or alertness were also used (e.g., Fernández-Juricic et al., 2009; Kaby & Lind, 2003; Lima & Bednekoff, 1999; Rogers et al., 2004). These studies have found a period between these subtle behavioral shifts and overt escape actions (response time delay), which is considered a crucial risk-assessment phase (Cresswell et al., 2009). This phase has been found to depend on multiple factors such as species, context (e.g., food availability, surrounding environment: Fernández-Juricic et al., 2002), individual characteristics (Jones et al., 2009) and flock size (Boland, 2003). An issue remains, however, that even these subtle behaviors might not accurately capture the time when an animal detects an approaching threat (Barbosa & Castellanos, 2005; Fernández-Juricic, 2012; Lima & Dill, 1990; Tätte et al., 2019), due to the lack of a more direct measure of visual attention.

### Visual sensory ecology of birds

More recent research in visual sensory ecology has emphasized the role of vision in the context of vigilance. Vision is crucial for gathering distant predator cues, and it is thus expected to play a significant role in predator-prey interactions (Barbosa & Castellanos, 2005). Specifically, predation is considered one of the primary drivers of the evolution of the visual system in birds, as well as foraging (Martin, 2017). Birds’ retinal specializations, such as the *area* (a region of the bird’s retina with a high density of photoreceptors) or the fovea (a pit-like area in the retina with high concentration of cone cells where visual acuity is highest, and is responsible for sharp, detailed, and color vision (see Bringmann, 2019 for more details)), are thought to be crucial for the detection, identification and tracking of predators (Fernández-Juricic, 2012; Martin, 2017) as well as risk assessment and response selection (Cresswell, 1993; Cresswell et al., 2009; Fuchs et al., 2019; Veen et al., 2000). Experimental studies tracking the foveal projections demonstrated that birds indeed use their foveas to inspect predator cues (Butler & Fernández-Juricic, 2018; Tyrrell et al., 2014; Yorzinski & Platt, 2014). Given the diversity and complexity of birds’ visual field configurations, it is necessary to understand how different types of visual information are acquired based on body posture and head orientation, rather than simply to consider a head-up and head-down dichotomy (Fernández-Juricic et al., 2004). Notably, while it is a prevalent notion that vigilant individuals are quicker to detect predators, this fundamental assumption has seldom been directly scrutinized in the literature (Beauchamp, 2015).

### Experimental rationales

This study thus leveraged recent fine-scale tracking methods to re-evaluate how a vigilant individual utilizes its high-acuity vision (such as the fovea) to detect a predator cue and subsequently guide individual and collective escape responses in the context of foraging and vigilance. We utilized a state-of-the-art, non-invasive motion capture technique to monitor the head orientations of pigeons moving freely within a flock (Kano et al., 2022; Nagy et al., 2023; Fig 1a and b). Motion capture cameras track with high accuracy the 3D position of markers, which, when attached to the pigeon’s head and body, enables to reconstruct the rotations of the head and body in all directions. This approach enabled us to closely analyze the birds’ fine-scale looking behaviors, while maintaining their natural foraging, vigilance, and social interactions; before, during and after simulated predation events (Fig 1c).

**Figure 1:**
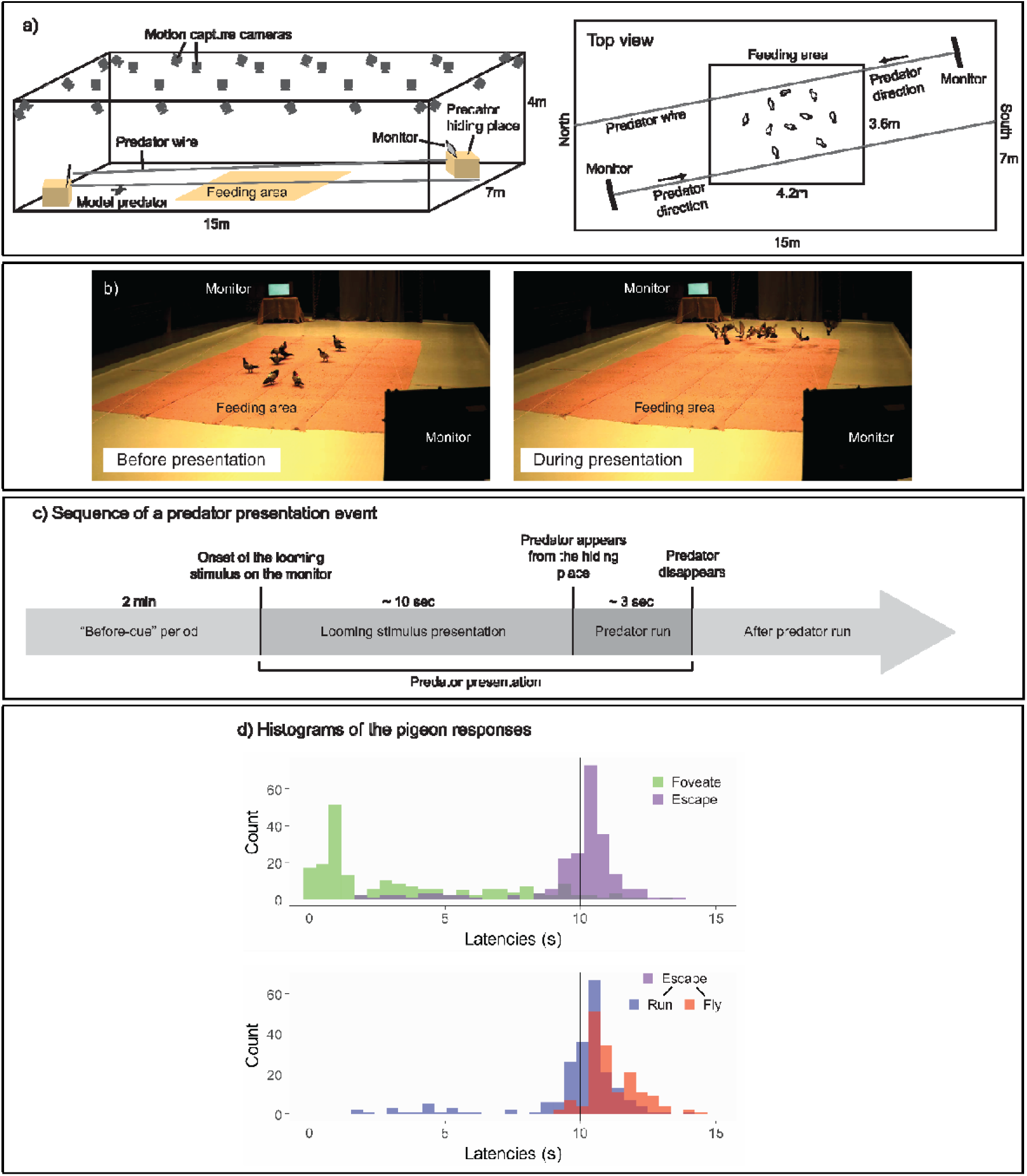
a) Experimental setup. In a large-scale motion capture system (SMART-BARN), the feeding area, the monitors and the predator running wire were installed (a 3D view on the left, and a top view on the right). b) Still frames captured from the periods before (top) and during (bottom) the predator presentation, respectively. c) Sequence of an event presenting a model predator. Each predator presentation event started with the presentation of a looming stimulus (predator silhouette), which lasted for approximately 10 s, on either of the two monitors, followed by the model predator running on the wire across the room (lasting for approximately 3 s). The “before-cue” period is defined as the 2 minutes directly preceding the onset of the looming stimulus. d) Histograms of the pigeons’ responses. Top: latency to foveate (green) and latency to escape –either running or flying (purple); bottom: latency to run (blue) and latency to fly (red) separately. The vertical line shows the looming stimulus offset.

Pigeons are well suited for our study system because they are relatively well studied for their visual system. In brief, although a relatively high acuity is maintained over the retina (Hayes et al., 1987), they have one fovea centrally located in the retina of each eye, with an acuity of 12.6 c/deg (Hodos et al., 1985). Their fovea projects laterally at ∼75° into the horizon in their visual field. They mainly use their foveas to attend to objects or conspecifics in the distance (>0.5m; Kano et al., 2022; Nalbach et al., 1990). Pigeons have another sensitive spot in their retina, the red field, which projects to the ground in their visual field. They mainly use the red fields in a foraging context, to search and peck seed on the ground in a close distance (<0.5m; Kano et al., 2022; Nalbach et al., 1990). Pigeons have a binocular overlap in the front covering an angle of approximately 20°, which they mainly use for pecking, perching, or attending to slow moving objects (Bloch et al., 1984; Green et al., 1992). Their blind area covers an angle of approximately 40° at the back of the head (Hayes et al., 1987; Martin, 1984). Due to these systematic structures, we assumed that their head movement indicates critical information about vigilance, foraging, and detection. Yet, it should be noted that their eye movement was not tracked in our system, although it is typically confined within a 5 degrees range (Wohlschläger et al., 1993). We thus considered this estimation error of the foveation (directing visual focus to the fovea to achieve the clearest vision) in our analysis, as a part of the error margin (see Methods).

In our observations, pigeons foveated on the predator cue typically before the offset of looming stimulus (Fig 1d); we considered this foveation as a potential indicator of early detection. Pigeons initiated running and then took flight typically during a predator presentation event (Fig 1d, see also Table S1). Pigeons responded either directly to the looming stimulus or after the appearance of the model predator, by running away and/or flying. Therefore, our first measure of escape was the presence of an escape response (either running or flying) prior to the looming stimulus offset (i.e., whether the pigeon runs or flies before the looming stimulus disappears). Our second measure of evasion focused on flight responses (predominantly occurring after the predator appeared), given that flight is the most prominent form of escape response in birds and is widely noted in the literature for its contagious nature among flockmates (Davis, 1975; Lima, 1995b; Lima & Zollner, 1996).

### Experimental hypotheses

We subsequently developed a series of hypotheses outlined in Table 1. The initial two hypotheses (Hypotheses 1 and 2) aim to examine whether foveation correlates with predator detection. Hypothesis 1 posits that pigeons employ their foveas to evaluate the predator (looming) cue and Hypothesis 2 that the foveation latency is correlated with vigilance-related behaviors of the individual. Hypothesis 2 was further divided into 2 sub-hypotheses; respectively assessing if the foveation latency depends on the state of the individual at the onset of the looming cue, and depends on the general behavior of the pigeons before the cue appears. Specifically, Hypothesis 2.1 suggests that the birds’ behavioral state observed at the onset of the looming stimulus (e.g., head-up, foraging) predicts the latency to foveate on the cue. Additionally, we assessed spatial positioning—relative to the threat (distance from the monitor) or to conspecifics (nearest neighbor)—as these factors are known to influence vigilance and escape behaviors (Beauchamp, 2008; Beauchamp & Ruxton, 2008; Inglis & Lazarus, 1981). Hypothesis 2.2, posits that various aspects of vigilance/foraging behaviors observed prior to the presentation of the looming stimulus (e.g., head-up, pecking rate, head-saccade rate, time spent foveating on predator-related objects or conspecifics) correlate with the latency to foveate on the cue. As noted, these related yet distinct facets of visual/foraging behaviors have been identified as differentially related to vigilance and escape (Cresswell et al., 2003; Jones et al., 2007, 2009). From Hypothesis 3, we asked whether earlier foveation on the predator cue predicts quicker escape responses by individual birds. While this is a common assumption in the literature, several factors appear to affect the length of the response time delay (Cresswell et al., 2009; Jones et al., 2009; Tätte et al., 2019). The final hypothesis (Hypotheses 4) focuses on the collective dynamics of detection and escape. Specifically, Hypothesis 4.1 posits that the first pigeon in a flock to foveate on the looming stimulus facilitates the escape behaviors of other members. A related hypothesis, Hypothesis 4.2, posits that the timing of escape initiations is socially contagious, specifically that the escape responses within a predator presentation event are more clustered than would be expected by chance.

**Table 1.** A set of hypotheses testing the assumptions about vigilance.

|  | Hypothesis | Test | Response | Description of the test |
| --- | --- | --- | --- | --- |
| 1 | Pigeons foveate on the predator cue (monitor or predator hiding table) during the 10-sec presentation period of the looming stimulus. | Linear Mixed Model: LMM | Proportion of time foveating on the objects of interest | Testing the effect of: Object type (the monitor presenting the looming stimulus, the monitor that did not present the looming stimulus, and the nearest neighbor pigeon) |
| 2.1 | Pigeons that are head-up (while not grooming nor courting or courted) foveate on the predator cue earlier than those that are feeding at the onset of the looming stimulus. | LMM | Latency to foveate on the predator cue. | Testing the effects of: Behavior exhibited at the onset of the looming stimulus (Head-up/Feeding), Distance from the monitor presenting the looming stimulus, and Distance from the nearest neighbor conspecific. |
| 2.2 | Pigeons' vigilance/feeding-related behaviors during the 2-min "before-cue" period predict their earlier foveation on the predator cue. | LMM | Latency to foveate on the predator cue. | Testing the effects of: Vigilance/feeding-related behaviors (proportion of time spent head-up, saccade rate, pecking rate, proportion of time foveating on any monitor, proportion of time foveating on any conspecifics). |
| 3 | Pigeons' first foveation on the predator cue predicts their escape. | LMM and Generalized Linear Mixed Model: GLMM | Probability of evasion before the offset of the looming stimulus, latency to fly. | Testing the effect of: Latency to foveate on the predator cue. |
| 4.1 | The first pigeon foveating on the predator (looming) within a flock facilitates the escape behaviors of the other members. | LMM and GLMM | Probability of escape before the offset of the looming stimulus and latency to fly in the other members (excluding the first pigeon). | Testing the effect of: Latency to foveate on the predator cue by the first pigeon. |
| 4.2 | Escape responses within a predator presentation event are more clustered than expected by chance. | Permutation test | Time interval between the successive escape (or flying) responses of pigeons within a given predator presentation event. | Comparing the mean interval observed in the data with the mean intervals expected by chance in a control event created by shuffling the latencies between event ID. |

## Results

### Overview of observations

We tested 20 pigeons across 6 trials each, with every trial involving a flock of 10 pigeons and consisting of 2 predator presentation events. This design resulted in a total of 240 observations, with each observation representing data from a single event for an individual pigeon. Each model incorporated this number of observations while excluding null cases in which pigeons displayed no response or cases where the system lost track of the pigeons (see Method for the details).

Within a few predator presentation events, our pigeons rapidly decreased their latencies to foveate, escape, and fly, indicating that learning occurred rapidly during these trials (Fig S1). At the very first predator presentation event, approximately half of the flock members did not foveate on the monitor nor escaped even after the appearance of the model predator (Table S1). From the second predator presentation event, most pigeons made foveation, escape, and flight responses. Our pigeons maintained these responses throughout the trials, although in the last few trials pigeons decreased their flight responses, indicating that habituation was minimal.

### Use of foveas during predator cue inspection (related to Hypothesis 1)

From the markers coordinates, we were able to reconstruct for each pigeon their head orientations simultaneously (Fig 2a-b). In the heatmaps (Fig 2c), we projected the 3D representations of the three objects of interest—the monitor displaying the looming stimulus, the monitor without the cue serving as a control, and the nearest conspecific (see supplementary for the 3D object definitions)—onto the pigeons’ local head coordinate system. We sampled this data during the period of the looming stimulus presentation and before any escape response by the pigeon (normalized to unit sum). Visual inspection suggested that pigeons foveated on the monitor displaying the looming stimulus, more so than on the other monitor that was not presenting the looming stimulus or on the nearest conspecific. It should be noted that the two monitors were placed on opposite sides of the room. Consequently, a pigeon located in the center and foveating on one monitor would have the other monitor falling ∼105° in azimuth angle on the opposite side of its head. This is likely resulting in the spot next to the foveal region for the monitor not displaying the looming stimulus (see Fig 2c).

**Figure 2:**
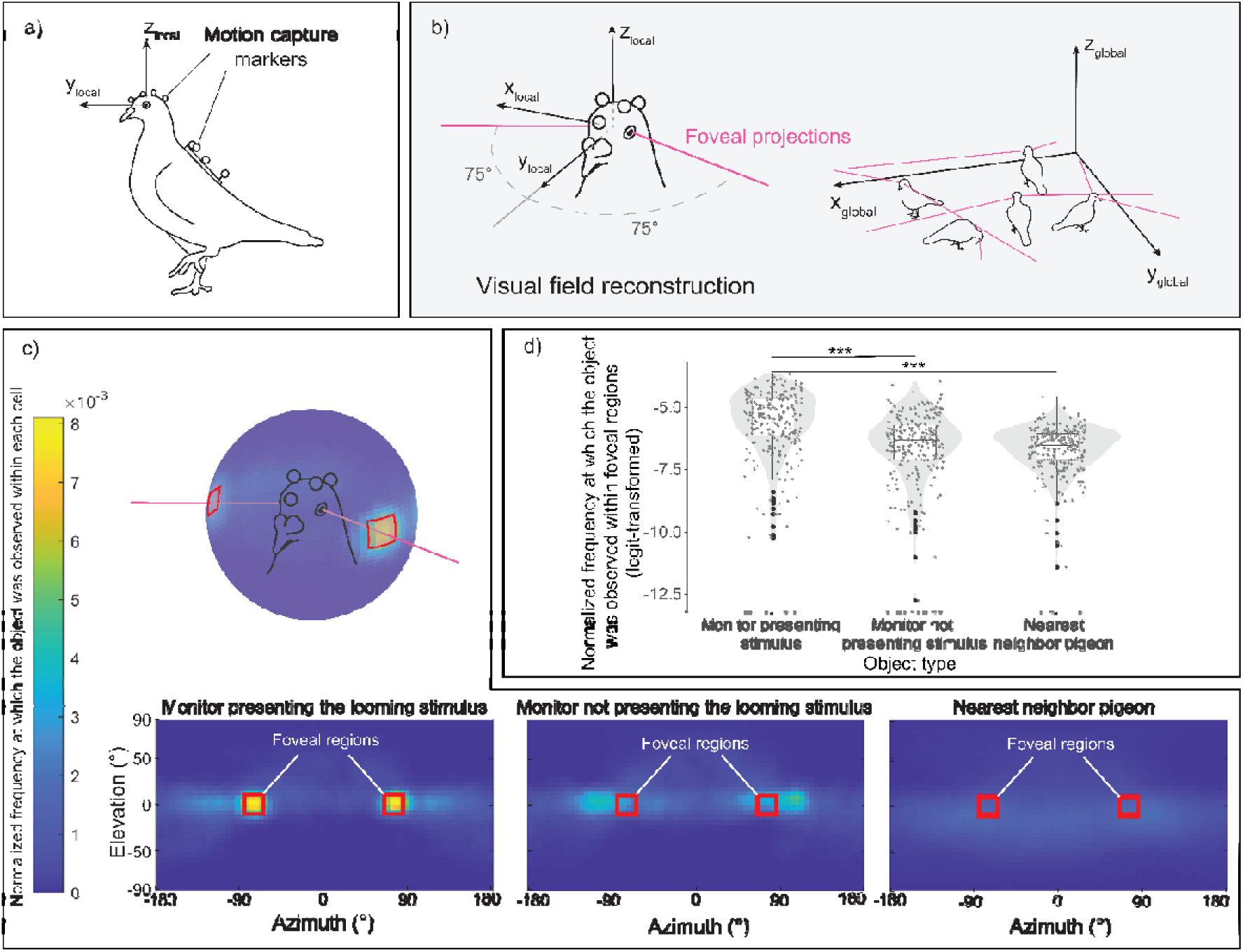
Pigeons’ foveation during the looming stimulus presentation. a) A pigeon equipped with motion capture markers on its head and back. b) Reconstruction of estimated foveal projections in the local coordinate system of the head (left) and in the global (motion-capture) coordinate system (right). c) Reprojection of each object of interest—the monitor presenting the looming stimulus, the monitor without the cue acting as a control, and the nearest conspecific—within the visual field of all pigeons across all trials. The red squares denote the designated region of estimated foveal projections in the visual field, defined as 75±10° in azimuth and 0±10° in elevation. The color scale illustrates the normalized frequency (normalized to a unit sum) at which each object of interest was observed within each 5×5° cell of the heatmap. d) Normalized frequency (normalized to unit sum then logit-transformed) at which each object of interest was observed within the defined foveal regions on the heatmap. The dots represent single observations (jittered horizontally for visualization), the grey shade represents the violin plot of the distributions, and the boxplot’s boxes represent the 0.25 and 0.75 quartiles (with the median represented as a line inside the box) and the whiskers the minimum and maximum values within the lower/upper quartile ± 1.5 times the interquartile range.

When testing for the normalized frequency at which each object was observed within the foveal region, we observed a significant effect of the object type ( ^2^(1) = 108.06, p < 0.001) (Fig 2d). Subsequent follow-up analyses revealed that pigeons focused on the monitor displaying the looming stimulus for a longer period compared to the other two objects (vs. monitor not presenting the looming stimulus, _χ_^2^(1) = 62.44, p < 0.001; vs. nearest neighbor, _χ_^2^(1) = 97.41, p < 0.001).

### Testing the assumptions about vigilance (related to Hypothesis 2)

The analysis predicting the latency to foveate with behavioral state (either head-up or feeding) and spatial factors (the distance from the monitor and from the nearest neighbor) at the onset of the looming cue reveal that it was significantly influenced by the behavioral state (χ^2^(1) = 13.78, p < 0.001) (Fig 3a). However, it was not significantly affected by the other spatial factors, such as the distance from the monitor (χ ^2^(1) = 1.20, p = 0.27) or the distance from the nearest neighbor (χ^2^(1) = 0.05, p = 0.82).

**Figure 3:**
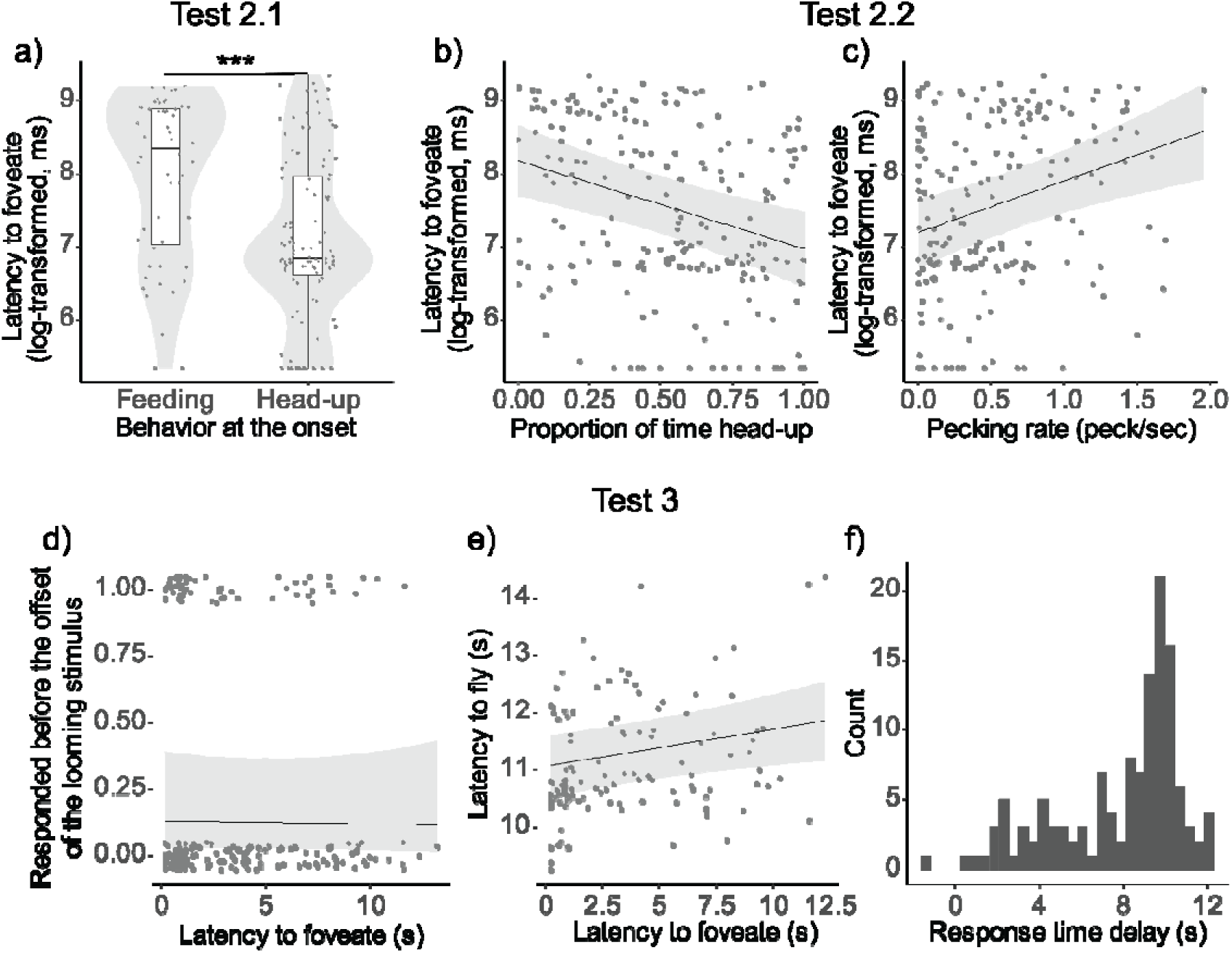
Results of the models from Test 2 and 3. a) Latency to foveate on the predator cue (log-transformed) as a function of behavioral state (feeding or head-up) at the onset of the looming stimulus. The dots represent single observations (jittered horizontally for visualization), the grey shade represents the violin plot of the distributions, and the boxplot’s boxes represent the 0.25 and 0.75 quartiles (with the median represented as a line inside the box) and the whiskers the minimum and maximum values within the lower/upper quartile ± 1.5 times the interquartile range. b-c) Latency to foveate on the predator cue as a function of the proportion of time spent being head-up during the “before-cue” period (b) and as a function of the pecking rate during the same period (c). The graphs for the 3 other variables can be found in Figure S6. d-e) Probability of escaping before the looming stimulus offset (d), and latency to fly (e) as a function of the latency to foveate. (f) Histogram of the distribution of the response delay (time between the latency to foveate and the latency to fly). For all depicted results, regression lines were determined with other variables held constant, set to their mean values. A comprehensive table detailing the outcomes of the models can be found in Table S7.

In addition, the latency to foveate was significantly predicted by most of the feeding-/vigilance-related behaviors of the pigeon during the “before-cue” period, including the proportion of time spent being head-up (χ ^2^(1) = 15.90, p < 0.0001), the pecking rate ( ^2^(1) = 14.07, p = 0.0002), the proportion of time foveating on a monitor (χ ^2^(1) = 12.04, p = 0.0005), and the saccade rate ( ^2^(1) = 9.85, p = 0.0017). However, the proportion of time foveating on any conspecifics did not have a significant effect (χ^2^(1) = 0.06, p = 0.8116) (Fig 3b-c, Fig S6).

When comparing the Akaike Information Criterion (AIC) of all five models, the most effective model included the proportion of time spent being head-up (AIC = 609.99). This was followed by models including the pecking rate (AIC = 612.40), the proportion of time foveating on the monitor (AIC = 614.01), and the saccade rate (AIC = 616.00).

In further analyses outlined in the Supplementary Material, we examined the changes in these vigilance- and feeding-related behaviors shortly after the predator disappeared (1 minute after the predator’s disappearance) as compared to the 1 min period preceding the stimulus onset. The results highlighted a significant increase in vigilance and a significant decrease in feeding immediately following the predator’s disappearance across all variables (see Fig S5 and Table S5 for details).

### Detection and escape (related to Hypothesis 3)

The model predicting the probability to escape before the looming stimulus offset indicated that the latency to foveate had no significant effect ( ^2^(1) = 0.02, p = 0.8716). However, the model predicting the latency to fly revealed a significant effect of the latency to foveate, suggesting that earlier foveation predicted quicker flight responses ( ^2^(1) = 6.49, p = 0.0108) (Fig 3d-e). We observed that the response time delay (the interval between the latency to foveate and the latency to fly) was relatively lengthy and exhibited considerable variation (Fig 3f). During this period, pigeons rarely returned to the feeding activity, as indicated by the low mean pecking rate (0.10 pecks/sec). In the Supplemental Material, we demonstrate that this variation can be partially attributed to individual differences; a within-individual repeatability analysis revealed that one contributing factor to this variation is inter-individual differences (R = 0.128, p = 0.0404).

### Social contagion of escape responses (related to Hypothesis 4)

Earlier foveation of the first pigeon was not significantly related to an earlier escape responses among the other flock members, although there was a trend ( ^2^(1) = 3.66, p = 0.0559). We then examined the same model with latency to fly as a continuous response and found that the earlier foveation of the first pigeon significantly predicted earlier flying responses among the other flock members ( ^2^(1) = 17.32, p = 0.0003) (Fig 4a-b).

**Figure 4:**
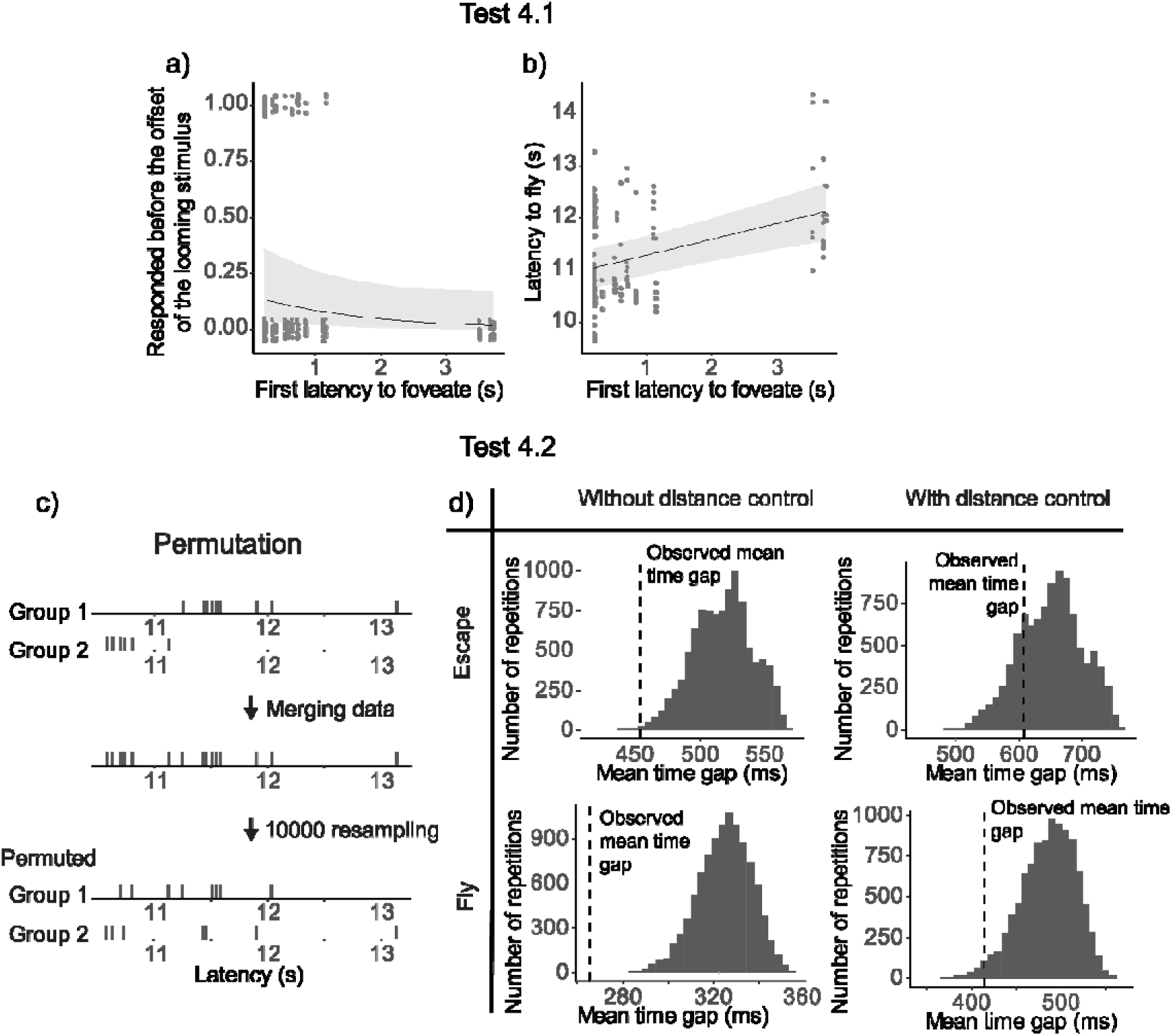
Results of test 4. a-b) Latency to foveate of the first individual foveating on the monitor as a function of the other individuals’ probability of escape before the cue offset (a) and the other individuals’ latency to fly (b). All the regression lines were calculated with other variables constant and equal to their mean value. A detailed table of the models’ results is available in Table S7. c) An overview of the permutation test procedure. d) Results of the permutation test. Histograms denote the distributions of the average time gap in permutated data. The vertical dashed line indicates the mean time gap from the observed data. The proportion of permutated data less than the observed data (on the left of the dashed line) gives the p-value; escaping (top) and flying (bottom); without (left) or with (right) distance control.

To test Hypothesis 4.2, we conducted a permutation test to check if the time gaps between individual departures were significantly more clustered than expected by chance. In short, we merged departures latencies from 2 different events, then resampled the latencies to obtain an “expected mean time gap” between 2 individual latencies (Fig 4c, see Methods for more details). The mean time gaps from the observed data were then compared to those from the 10,000 permutations to determine a p-value. The observed time gaps were significantly shorter than the permuted gaps for both escape (p = 0.00038) and fly responses (p < 0.0001; Fig 4d).

Upon inspecting the data, we found a potential confound effect of the distance of the flock from the monitor on the latency to escape or fly, and therefore we ran a second permutation test controlling for the distance (see Methods for more details). Using these controlled datasets, we found that the mean time gap was not significantly different for the escape latency (p = 0.206), but it was significantly shorter for the fly latency (p = 0.0237) (Fig 4d).

## Discussion

This study leveraged fine-scale behavior tracking to examine key assumptions of vigilance research in pigeon flocks during free foraging, particularly focusing on the role of foveation. Using motion-capture technology, we tracked the individuals’ visual fields and quantified their behaviors automatically during predator presentation events. Our findings supported the assumptions about vigilance in two aspects: foveation in pigeons is associated with predator inspection, and behaviors related to vigilance and feeding are reliable predictors of earlier foveation in pigeons. Our findings also largely supported the assumptions about collective vigilance in two aspects: earlier foveation is indicative of earlier flight responses at both individual and collective levels, and there is a social contagion effect in their evasion flight responses. However, these results are somewhat complicated by the individuals exhibiting long and variable response time delays—the interval between foveation and escape responses. Moreover, social contagion of evasion was only observed in flight responses following the appearance of the model predator (after most flock members had foveated on the predator cue), not in earlier responses. In summary, while our results largely affirm the prior assumptions about vigilance, we have identified several confounding factors, which will be discussed further below.

### Use of foveas during predator cue inspection (related to Hypothesis 1)

Our pigeons primarily use their foveas to inspect the looming stimulus. This is consistent with previous studies showing that starlings and peacocks use their foveas for predator inspection (Butler & Fernández-Juricic, 2018; Tyrrell et al., 2014; Yorzinski & Platt, 2014). Our research extends these findings to situations where pigeons forage freely on the ground in a flock. It has been previously shown that pigeons use their frontal visual fields for searching and pecking at grain, and their foveas to inspect distantly presented objects (Kano et al., 2022). In line with this, our study found that pigeons did not employ their frontal visual fields to inspect the looming stimulus. Additionally, pigeons did not use their foveas to view conspecifics during the stimulus presentation. Collectively, our results indicate that, during the stimulus presentation, pigeons primarily relied on their foveas to attend to the predator cue. This consolidates the idea that the avian fovea is crucial in predator detection and inspection (Martin, 2017).

### Testing the assumptions about vigilance (related to Hypothesis 2)

We found that vigilance-related behaviors, including keeping the head-up, engaging in more frequent head movements (saccades), and earlier monitoring of potential threat locations (i.e., monitors), indeed lead to earlier foveation on the predator cue. Conversely, feeding-related behaviors, such as keeping the head-down and a higher pecking rate, delay the foveation on the cue. The spatial configuration of the flock, such as the distance from the predator cue and foveation on conspecifics, did not influence earlier foveation on the predator cue. Further analyses showed that these vigilance-related behaviors significantly increased, and these feeding-related behaviors decreased after the presentation of the predator (see Fig S5).

These findings indicate that all the measured vigilance- and feeding-related behaviors are relevant to predator detection in our study system. This is in line with several studies (Devereux et al., 2006; Hilton et al., 1999; Lima & Bednekoff, 1999), though not others (Cresswell et al., 2003; Jones et al., 2007, 2009; Kaby & Lind, 2003). While prior research has suggested that species with a relatively large visual field can maintain vigilance peripherally (Fernández-Juricic et al., 2004), our data propose that behaviors such as head-up and scanning are still advantageous, and head-down while feeding is still costly to predator detection, even in pigeons that possess a relatively large visual field. Furthermore, previous studies have noted that, in blue tits, vigilance- and feeding-related behaviors are linked to detection only when the feeding task is more demanding (Kaby & Lind, 2003). This might be partly relevant to our results, as we used a moderately attention-demanding task where pigeons searched for grain in a grain-grit mixture.

Most notably, our fine-scale tracking of pigeon behaviors may have refined our ability to identify predator detection. Previous studies have inferred predator detection by using escape responses as the sole measure (Kenward, 1978; Lima & Zollner, 1996; Quinn & Cresswell, 2005) or by observing more subtle signs of alertness such as freezing, adopting a straight upright posture, crouching, or ceasing to feed (e.g., Fernández-Juricic et al., 2009; Kaby & Lind, 2003; Lima & Bednekoff, 1999; Rogers et al., 2004), or by combining these responses (Cresswell et al., 2003; Lima, 1995a, 1995b; Tisdale & Fernández-Juricic, 2009; Whittingham et al., 2004). However, as previously claimed (Barbosa & Castellanos, 2005; Fernández-Juricic, 2012; Lima & Dill, 1990; Tätte et al., 2019), predator detection might occur earlier without overt behavioral responses, which can be too subtle to detect under standard observational conditions. In our study, we measured the birds’ first foveation on the predator cue while excluding instances of short and early foveation on the cue (see Method). Although making definitive claims is technically challenging without assessing the internal perceptual process, the clear relationships observed between foveation and vigilance- and feeding-related behaviors in our results suggest that foveation is likely a reliable proxy for “detection” within our study system.

### Detection and escape (related to Hypothesis 3)

In our study, earlier foveation on the predator cue predicted earlier evasion responses from the model predator. However, this relationship was confirmed only when we analyzed the flight responses made after the predator’s appearance, not when we included pigeons’ running as the escape responses. Moreover, pigeons exhibited relatively long and variable response times between foveation and the flight response. This response time delay was, in part, related to consistent individual differences across trials.

One interpretation of this result is that pigeons, after foveating on the predator cue, assessed its potential risk. The influence of earlier detection on the decision to fly earlier likely stems from increased sensitivity to risk following the appearance of the model predator. The consistent individual differences in response time delay suggest variations in risk sensitivities among individuals. The relatively long and variable response time delay observed is likely inherent to our study design, which involved using a looming stimulus as a warning for the appearance of a model predator. Notably, pigeons rapidly decreased their latency to foveate after just a few presentations (as shown in Fig S1), indicating increased sensitivity rather than habituation to the stimuli across repeated presentations. The low pecking rate during the response time delay (as detailed in the Results section) suggest that pigeons were alert when viewing the looming stimulus. However, this alertness more likely led them to assess the stimuli rather than to initiate an explicit escape response in most cases. These findings are in line with the previously established idea that detection and escape represent two distinct stages in antipredator responses (Barbosa & Castellanos, 2005; Fernández-Juricic, 2012; Lima & Dill, 1990), and that response time delay is associated with risk assessment (Cresswell et al., 2009) and individual differences (Jones & Godin, 2010).

### Social contagion of escape responses (related to Hypothesis 4)

When the first pigeon in the flock foveated on the predator cue earlier, the remaining flock members flew earlier. Permutation tests indicated that their evasion flight responses are socially contagious (no clear evidence for earlier escape response involving running).

One interpretation of these results is that the decision to fly by the first pigeon that foveated triggered a following response from the rest of the flock. This social contagion in flight responses has been previously demonstrated in birds, including pigeons (Davis, 1975). Notably, our pigeons did not directly foveate on other flock members (as illustrated in Fig 2), suggesting that direct foveation on other pigeons is not necessary for the social contagion of flights. This finding is consistent with the notion that birds only need to maintain peripheral visual contact with their flock members to observe and respond to their departure flights (Lima & Zollner, 1996). Additionally, considering the findings related to Hypotheses 3 and 4, our results indicate a nuanced form of collective detection: while earlier detectors may trigger earlier flight responses in the flock via social contagion, non-detectors do not necessarily gain an advantage from these detectors, as most flock members likely already detected the predator cue.

### Limitation

A potential limitation of our study is its limited generalizability. We focused on pigeons, which are middle-sized birds, and their escape responses may differ from those of smaller birds commonly studied in vigilance research; pigeons might, for instance, be more hesitant to take flight due to the higher energy costs related to their weight. Furthermore, while our system precisely tracked freely foraging birds, some elements of our experimental setup were artificial. The looming stimulus and the model predator were presented in a somewhat disjointed manner. Pigeons had to rapidly learn the association between these two stimuli over several trials, potentially increasing their reluctance to fly. This learning process and the disjointed stimulus presentation might have contributed to the long and variable response time delays observed in our study. While these delays provided a unique insight into collective escape, particularly in cases where early detection does not immediately result in early escape, it is possible that this scenario might be more likely with a more naturalistic predator stimulus. Future research could address these limitations by conducting similar studies in the field, outside of the motion-capture system, perhaps using the emerging technology of markerless 3D posture tracking in birds (Waldmann et al., 2022, 2023).

### Conclusion

Our research largely supports the common assumptions in vigilance studies. Key among our findings is the relationship between foveation and predator inspection, along with vigilance- and feeding-related behaviors. This suggests that foveation can be a useful proxy for detection in bird vigilance studies. Additionally, we observed that earlier foveation led to earlier flight responses in the flock, facilitated by social contagion. Interestingly, this contagion occurred even though most flock members likely detected the predator cues, as suggested by the long and variable response time delay, which are likely associated with risk assessment. Therefore, our results imply that collective escape does not always involve non-detectors benefiting from detectors. Our study highlights the importance of considering the vision as well as the disparity between detection and escape responses in future vigilance research.

## Methods

### Subjects

Twenty pigeons (*Columba livia*, 13 females and 7 males) participated in this study (492 ± 41g; mean ± SD). All pigeons originated from a breeder and were juveniles of the same age (1 year old). They were housed together in an aviary (2w x 2d x 2h m; w-width, d-depth, h-height) with perching and nesting structures. The pigeons were fed grains once a day. On experimental days, they were fed only after the experiments was completed; this ensures 24-hour no feeding at the time of the experiment, although we did not control the amount of the food over the course of the experimental periods. Water and grit were available in the aviary *ad libidum*.

### Ethics statement

All the experiments using animals in this study were performed under the license 35-9185.81/G-19/107 awarded by the Regierungspräsidium Freiburg, Abteilung Landwirtschaft, Ländlicher Raum, Veterinär-und Lebensmittelwesen, animal ethics authorities of Baden-Württemberg, Germany. Handling was reduced to a minimum to avoid stress. No animal was hurt or killed during the predator presentation tests, and they were transferred back to their home aviary directly after the tests. The pigeons were provided with perches and nesting structures in their aviary, and their health states were checked on a daily basis. After all experiments were performed, all the individuals were given to a breeder to continue living as breeding pigeons.

### Experimental setup

The experiment took place in SMART-BARN, the state-of-the-art animal tracking system build at the Max Planck Institute of Animal Behavior (Fig 1a; Nagy et al., 2023). We used the motion-capture feature of SMART-BARN, which was equipped with 32 motion-capture cameras in an area of 15w x 7d x 4h m (12 Vero v2.2 and 20 Vantage 5, VICON). At two opposite corners of the room, we placed tables covered with fabric (“predator hiding place”; 105w x 75d x 75h cm), which hide from the pigeons’ view a model predator (plastic sparrow hawk; wingspan 60cm and beak-tail length 35cm). At the center of the room, we placed jute fabric (4.2l x 3.6w m; l-length) where we scatted food to encourage the pigeons to stay there during the experiments (“feeding area”). The model predator ran on a thin wire across the room via a motored pulley mechanism (Wiral LITE kit, Wiral Technologies AS.) until it disappeared into another hiding place set at the end of the room. On top of each predator hiding table, we installed a monitor (61.5w x 37h cm, WQHD (2560 x 1440), 144Hz, G-MASTER GB2760QSU, Iiyama), which displayed a looming stimulus before the model predator appears from the hiding place. We specifically chose a monitor with high temporal resolution to match the pigeon’s Critical Flicker Fusion Frequency (threshold at which a flickering light is perceived by the eye as steady) that reaches up to 143Hz (Dodt & Wirth, 1954).

This looming stimulus was a silhouette of a predator mimicking the approach of a predator toward the pigeon flock. The looming stimulus lasted for 10 seconds, starting from a still small silhouette of a predator (20w x 4h cm) in the first 5 seconds and then expanding until the silhouette covers the screen entirely in the last 5 seconds. A looming stimulus was chosen because it is generally perceived as threatening across species (Evans et al., 2019), and is subtle enough so that pigeons require effort to detect it in our experimental setup. The looming stimulus was played back on a laptop PC (ThinkPad P17 Gen 2, Lenovo) connected to both monitors, and its onset was manually triggered by the experimenter. This onset time was recorded in the motion-capture system via analogue electric signal, specifically by the experimenter pressing a button at the onset of the looming stimulus on a custom Arduino device (Arduino UNO R3, Arduino) connected to the motion capture system. The same Arduino device also indicated the onset of the model predator to the experimenter via a small LED flash.

### Experimental procedures

#### General experimental design

Each pigeon underwent maximum one trial per day and a total of 6 trials. A flock of 10 pigeons participated in each trial, yielding 2 separate flocks in each trial, and a total of 12 trials in the whole experiment. Each pigeon was pseudo-randomly assigned into either flock in each trial, in such a way that each pigeon was paired within a group with any other pigeon at least once across all 6 trials. Each trial composed of 2 predator presentation events, and each event presents a model predator from either the hiding place (“North” or “South” side), yielding 24 events in the whole experiments. The order of presenting the North and South side of hiding places was counterbalanced across groups within each trial.

#### Preparation for the experiment

Before the experiment, the pigeons were transported from the aviary using a pigeon carrier (80w x 40d x 24h cm). They were then equipped with 4 motion capture markers (6.4mm diameter, OptiTrack) glued on the head feathers and 4 motion capture markers (9mm diameter) on a small solid styrofoam plate (7l x 3.5w cm); this small plate was attached to a backpack worn by each pigeon with Velcro. The experimenter then held the pigeon’s head briefly (less than a minute) in a custom 3D frame equipped with 4 web cameras and a triangular scale (26w x 29d x 27h cm) and then filmed the head with web cameras (C270 HD webcam, Logitech) synchronized by a commercial software (MultiCam Capture, Pinnacle Studio) in order to reconstruct the eyes and beak positions relative to the markers post hoc (see below for the reconstruction methods). Additionally, the motion capture system was calibrated just before the experiment starts with an Active Wand (VICON) until all cameras have recorded 2000 calibration frames.

#### Trial design

At the start of each trial, all 10 pigeons were released in the motion capture room. After 2 minutes of acclimatization period, an experimenter scattered a grain-grit mix evenly in the feeding area (Fig 1a-b). This grain-grit mix comprised of 200g of seeds with 400g of grit to make the foraging task moderately challenging. The experimenter then hid behind a curtain, outside the motion capture room. After the experimenter visually confirmed that minimally half of the pigeons (≥5 individuals) started feeding (pecking grain; this always happened within a few seconds after the experimenter started scattering the food), a free feeding period started. The duration of this period was randomized between 2 and 5 minutes across trials to reduce predictability of the predator event. The last 2 minutes of this feeding period, referred as the “before-cue” period, were used to extract pre-stimulus behavioral measures. At the end of the feeding period, the experimenter triggered the looming stimulus. After the offset of the looming stimulus (mean 10.83 ± SD 0.43 s after the onset), the experimenter ran the model predator on the wire, and the predator took approximately 3 seconds before it disappeared from the room (mean 2.83 ± SD 1.18 s; Fig 1c). After half of the pigeons resume feeding (within 1-4 min after the predator disappears), and another free feeding period of 2-5 minutes, a second predator presentation event occurred following the same procedures, except that the looming stimulus and the model predator appeared from the other monitor and hiding place. Each trial lasted for approximately 30 minutes. After the experiment, the motion-capture markers were detached from each pigeon before being brought back to the aviary.

## Data analysis

### Reconstruction of the pigeons’ head

A custom pipeline (see Data availability section) was used to reconstruct the relative 3D position of the center of the eyes, the beak-tip and the center of the markers from the 4 still images of the pigeon’s head (see above). These key points were manually labelled by an experimenter on each picture (an updated algorithm is now available for automatic labelling using YOLO, also included in the deposit). A custom MATLAB program based on structure-from-motion reconstructed the 3D coordinates of all the keypoints with a less than 2 pixels mean reprojection error. These procedures were confirmed to yield accurate reconstruction of head orientations, less than a degree of rotational errors (Itahara & Kano, 2022).

### Processing of the motion capture data

The motion capture data were exported as csv files from the motion-capture software (Nexus version 2.14, VICON). The csv files were then imported and processed in the custom codes written in MATLAB (provide in Kano et al., 2022, also provided in our deposit). The motion capture data consisted of a time series of 3D coordinates of the markers attached to the head and back of the pigeons (Fig 2a). From the reconstructed 3D positions of eye centers and beak tip, the local coordinate system of the pigeon’s head (the location of the objects and conspecifics from the pigeon’s perspective) was defined (Fig 2b). In this local coordinate system, the horizon (the elevation of the local X and Y axes in the global coordinate system) was 30° above from the principal axis of the beak, and the X, Y, Z axis pointed to the right, front, and top of the pigeon’s head, respectively. It should be noted that this angle of 30° was determined based on the typical standing postures of pigeons, as identified in Kano et al. (2022). While this previous study determined the angle on a trial-by-trial basis, our study employed a fixed angle following previous recommendations, and also due to the small variation of this angle across trials and individuals.

The data were filtered using the custom pipelines described in Fig S8 of Kano et al. (2022). Briefly, the raw motion-capture data were gap-filled in NEXUS, then smoothed using custom MATLAB codes. The translational and rotational movements of the reconstructed local coordinate system were further smoothed, and any improbable movements were removed using the same pipelines. As a result, the loss of head local coordinate system data amounted to 10.07±11.79% (mean ± SD) per individual in each trial, and the loss of back-marker centroids was 2.53±0.98% (mean ± SD). Although the loss of head data was nontrivial, visual confirmation showed that most loss occurred when pigeons were self-grooming (thus self-occluding their head markers from the motion capture cameras); i.e., when they likely were not attending to any object of interest. Head saccades were also removed from all foveation data due to the possibility that visual processing is inhibited during saccades in birds (Brooks & Holden, 1973). It should be noted that one deviation from the previous study in terms of filtering parameters was that the data smoothing was performed at 60Hz, rather than 30Hz, following previous recommendations.

### Reconstruction of gaze vectors

We then reconstructed the gaze (or “foveal”) vectors of each pigeon based on the known projected angles of the foveas, 75° in azimuth and 0° in elevation in the head local coordinate system (Nalbach et al., 1990). Pigeons use these gaze vectors primarily when attending to an object/conspecific in the middle to far distance (roughly >50 cm from the head center; Kano et al., 2022). Although pigeons have another sensitive region of retina, known as red field (Wortel et al., 1984), this region was not examined in this study because pigeons primarily use this retinal region to attend to objects in the close distance (roughly <50 cm), such as pecking and searching for grain (Kano et al., 2022).

### Behavioral classification

“Foveation” was defined when an object of interest (such as the monitor or another pigeon) fell within ±10° of the gaze vectors. This margin was estimated to accommodate well the eye movement, which was typically within 5° in pigeons (Wohlschläger et al., 1993). Head-saccades were defined as any head movement larger than 5°, that lasted for at least 50ms and faster than 60°/s, and fixations were defined as inter-saccadic intervals, based on the previous study (Kano et al., 2022). Our objects of interest included the monitors, the tables that hid the predator, and conspecifics. These objects were defined as sphere encompassing them (see Supplementary Material and Fig S3). In our observations, pigeons foveated on the predator cue (monitor or predator hiding table) typically before its offset (Fig 1d); in a few instances (9 out of 120 observations), pigeons foveated on the model predator after the looming stimulus had disappeared, but these cases were excluded from our analysis. To ensure that the first foveation of a pigeon on the predator cue was not a result of random gaze crossing, we excluded instances of foveation that were shorter than 300 ms, based on the typical intersaccadic interval for a pigeon, which typically ranges from 300 to 400 ms (Kano et al., 2022). Moreover, to remove cases where the gaze vector was on the predator cue at the onset of the looming stimulus, we excluded from the analysis the first 200 ms of looming stimulus period based on typical reaction times of a pigeon, which is usually longer than 200 ms (Blough, 1977).

In addition to the foveation, the pigeons’ behavior (e.g., head-up, grooming, feeding, running, flying) was classified automatically based on simple thresholds using the 3D postural data (Table 2; for more detailed explanations, see Table S3); those thresholds were verified by two human raters (see Supplementary Materials and Table S4).

**Table 2.**
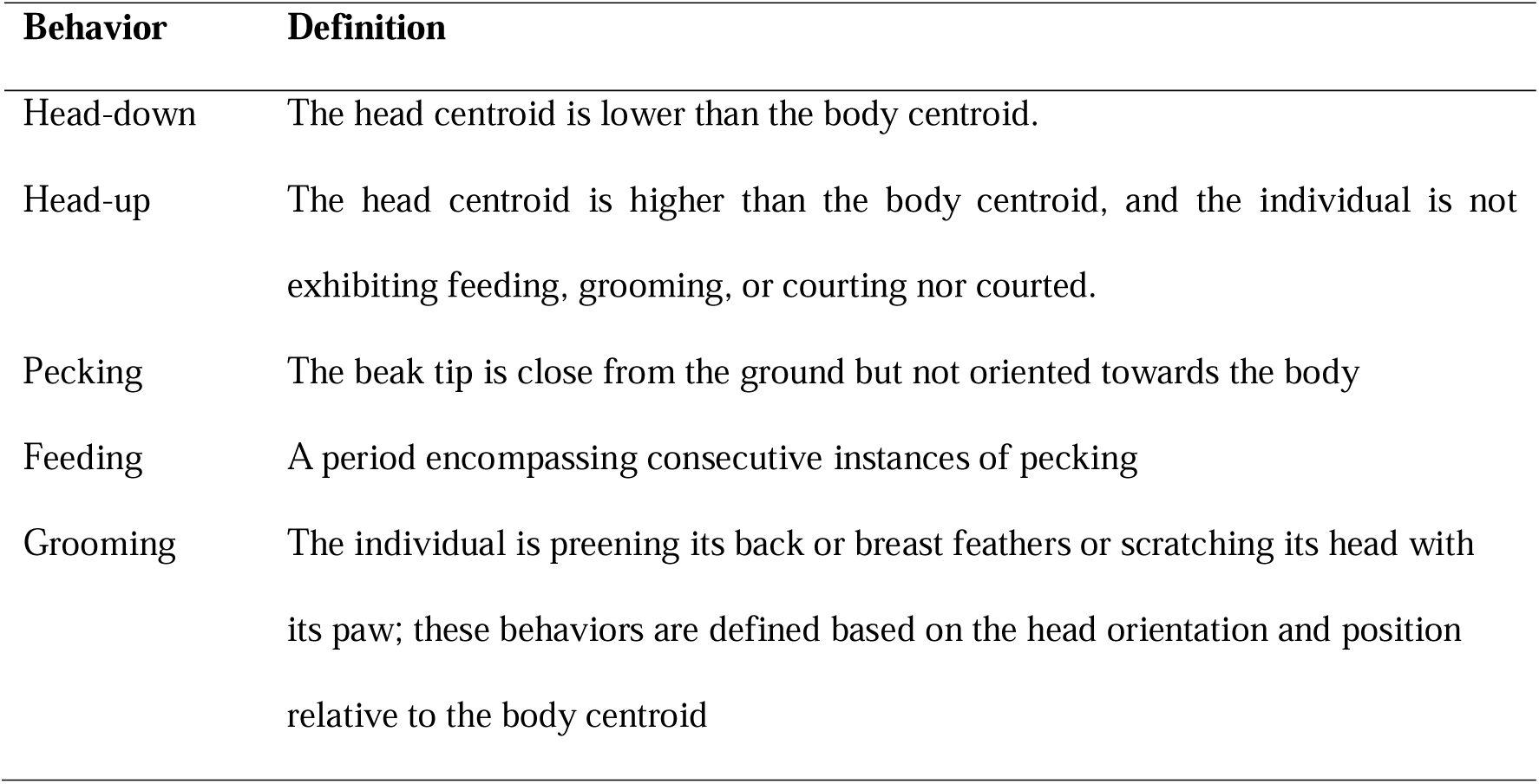
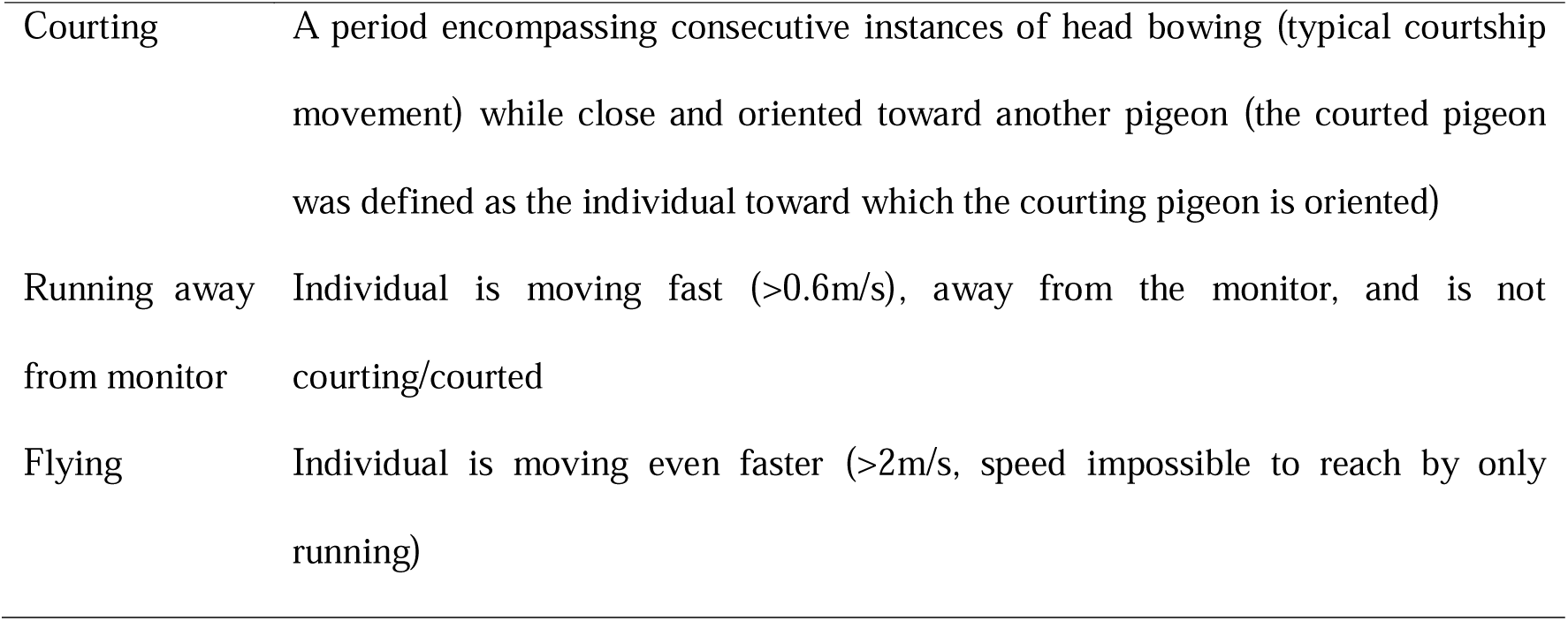
-definition of the automated classification of the behaviors. See Table S3.

| Behavior | Definition |
| --- | --- |
| Head-down | The head centroid is lower than the body centroid. |
| Head-up | The head centroid is higher than the body centroid, and the individual is not exhibiting feeding, grooming, or courting nor courted. |
| Pecking | The beak tip is close from the ground but not oriented towards the body |
| Feeding | A period encompassing consecutive instances of pecking |
| Grooming | The individual is preening its back or breast feathers or scratching its head with its paw; these behaviors are defined based on the head orientation and position relative to the body centroid |
| Courting | A period encompassing consecutive instances of head bowing (typical courtship movement) while close and oriented toward another pigeon (the courted pigeon was defined as the individual toward which the courting pigeon is oriented) |
| Running away from monitor | Individual is moving fast ($>0.6\text{m/s}$), away from the monitor, and is not courting/courted |
| Flying | Individual is moving even faster ($>2\text{m/s}$, speed impossible to reach by only running) |

### Statistical analysis

All the statistical analyses were performed in R (R Core Team, 2022). We mainly relied on linear mixed models or generalized linear mixed models (LMM or GLMM; lmer or glmer function from the lme4 package; Bates et al., 2015) unless otherwise mentioned, and the significance of the predictors was tested using likelihood ratio test. We verified the model assumptions by checking the distribution of the residuals in diagnostic plots (histogram of the residuals, qq-plot and plot of the residuals against fitted values). We transformed the response variable (logit- or log-transformation) when it improved the normality of the residual distribution. To test for collinearity, we also checked the variance inflation factor (VIF) of the predictors. In all models, the continuous predictors were normalized (z-transformed). For all models, we also included several control variables to ensure they did not confound with our test predictors: the event number within a trial (1 or 2), the predator side (“North” or “South”), and the sex of the subject, as well as the pigeon ID and the trial ID as random effects (see Table S6 for the used R formulas). We report the effects of test variables in the Result section and report the effects of all test and control variables in Table S7. Detailed descriptions about the responses and test variables can be found in Table S2.

Specifically, to test Hypothesis 1, we quantified the frequency (normalized to a unit sum) at which each object of interest was observed within the defined foveal regions on the heatmap (illustrated by red rectangles; corresponds to the fovea location, adjusted by ± 10 degrees in both elevation and azimuth). This quantification was conducted for each pigeon at every predator presentation event.

After logit-transforming this response to improve the normality of residuals, we conducted a Linear Mixed Model (LMM) predicting the normalized frequency at which an object was observed in the foveal region with the object of interest as a within-subject categorical test variable, in addition to control variables and random effects (see Table S7).

To test Hypothesis 2.1, we conducted a LMM using the latency to foveate (log-transformed to improve the normality of residuals) as the response variable. The pigeons’ behavioral state (either head-up or feeding) was included as a categorical test variable, while the distance from the monitor and the distance from the nearest neighbor were included as continuous test variables (in addition to the control variables and random effects; Table S7).

To test Hypothesis 2.2, we ran 5 LMMs, each of which include the latency to foveate as a response variable and any of the 5 related behaviors (the proportion of time spent head-up, the saccade rate, the pecking rate, the proportion of time foveating on the monitor and the proportion of time foveating on any conspecifics) as a continuous test variable (in addition to the control variables and the random effects). We further tested the relative predictive power of the different test variables by comparing the resulting models’ efficiency using AIC scores.

To test Hypothesis 3, we conducted a binomial Generalized Linear Mixed Model (GLMM) analysis using the probability of escape before the looming stimulus offset as a binary response variable and the latency to foveate as a continuous test variable (in addition to control variables and random effects). Subsequently, we analyzed a Linear Mixed Model (LMM) with the latency to fly as a continuous response variable, with the same test and control variables.

To test Hypothesis 4.1, we included into a GLMM the probability of escaping before the looming stimulus offset as a binary response and the first pigeon’s latency to foveate as a test variable (in addition to control variables and random effects). To test the effect on the flying responses, we ran a LMM with the latency to fly of the other pigeons as continuous response variable, with the same test and control variables.

To test Hypothesis 4.2, we conducted a permutation test based on time gaps between the latencies of individual pigeons to either escape or fly within a given event. This time gap was defined as the latency of the focal individual minus the latency of the immediately preceding individual. For the permuted data, we sampled one event from two different flocks of pigeons. For each sample, the model predator was presented to both flocks from the same side (North or South) during the same trial. We matched the trial and predator presentation side because, among the three control factors (trial, predator presentation side, and event), these two significantly influenced individual latencies as determined by a LMM (the event ID was not a significant factor in this confirmatory test). We combined the latency data from both events, randomly redistributed this data between the two events 10,000 times, and then recalculated the time gaps each time (Fig 4c). We then calculated whether the observed mean time gap was significantly smaller than the expected mean time gap from the permutation by calculating the proportion of permutation with an average time gap smaller than the observed one (corresponds to the p-value). We also considered the possibility that the individuals’ distance from the monitor could confound the observed effect. Specifically, if the two pigeon flocks occupied different locations across the two events, and the distance from the monitor influenced the individuals’ latency to escape or fly, the results might reflect spatial rather than social influences. Upon inspecting the data, this effect seemed plausible. To account for this potential confound, we determined the overlapping areas by calculating the range between the minimum and maximum distances of each flock and subsequently analyzed only the individuals within this overlapping area.

## Acknowledgements

We thank the members of the Max Planck Institute of Animal Behavior (MPI-AB) as well as the Centre for the Advanced Study of Collective Behaviour (CASCB) for their valuable supports in conducting this study. Special thanks are extended to Drs. Mate Nagy, Dora Biro, Oliver Deussen, and Iain D. Couzin, along with the members of MPI-AB and CASCB, for their insightful comments on our study. We also thank Drs. Inge Müller and Daniel Zuniga, as well as the caretakers, for their dedicated hosting and care of the pigeons. Finally, we thank Mathias Günther and Alex Chan for technical support. This study was financially supported by MPI-AB, the DFG Cluster of Excellence 2117 CASCB (ID: 422037984), and the CASCB BigChunk projects (ID: L21-07).

## Data availability

A OSF repository containing the code used for the processing and analysis can be found on the OSF website (Delacoux & Kano, 2023). Sample data is also provided for the user (the whole raw dataset could not be included for size reason). The repository contains the code for the automatic labelling of the head calibration of the pigeons, the Matlab pipeline used for the motion capture data processing and filtering, and the statistical analyses in R (along with the tables containing the processed data).

## Supplementary Materials

### Overview of observations

**Table S1:** Total number of individuals that exhibited foveation, escape, and flight behavior over the course of the trials and events. Each event was composed of a flock of ten pigeons.

| Trial | Event | Group | # individuals foveating | # individuals escaping | # individuals flying |
| --- | --- | --- | --- | --- | --- |
| 1 | 1 | 1 | 6 | 5 | 4 |
| 1 | 2 | 1 | 9 | 9 | 7 |
| 1 | 1 | 2 | 5 | 4 | 4 |
| 1 | 2 | 2 | 8 | 9 | 9 |
| 2 | 1 | 1 | 9 | 9 | 6 |
| 2 | 2 | 1 | 10 | 10 | 6 |
| 2 | 1 | 2 | 6 | 9 | 9 |
| 2 | 2 | 2 | 8 | 9 | 8 |
| 3 | 1 | 1 | 10 | 9 | 10 |
| 3 | 2 | 1 | 10 | 10 | 6 |
| 3 | 1 | 2 | 8 | 9 | 9 |
| 3 | 2 | 2 | 9 | 10 | 8 |
| 4 | 1 | 1 | 8 | 10 | 8 |
| 4 | 2 | 1 | 9 | 10 | 7 |
| 4 | 1 | 2 | 9 | 10 | 6 |
| 4 | 2 | 2 | 5 | 10 | 10 |
| 5 | 1 | 1 | 9 | 9 | 6 |
| 5 | 2 | 1 | 10 | 10 | 6 |
| 5 | 1 | 2 | 10 | 8 | 0 |
| 5 | 2 | 2 | 9 | 9 | 6 |
| 6 | 1 | 1 | 8 | 10 | 7 |
| 6 | 2 | 1 | 10 | 6 | 0 |
| 6 | 1 | 2 | 8 | 10 | 6 |
| 6 | 2 | 2 | 8 | 10 | 9 |

**Figure S1:**
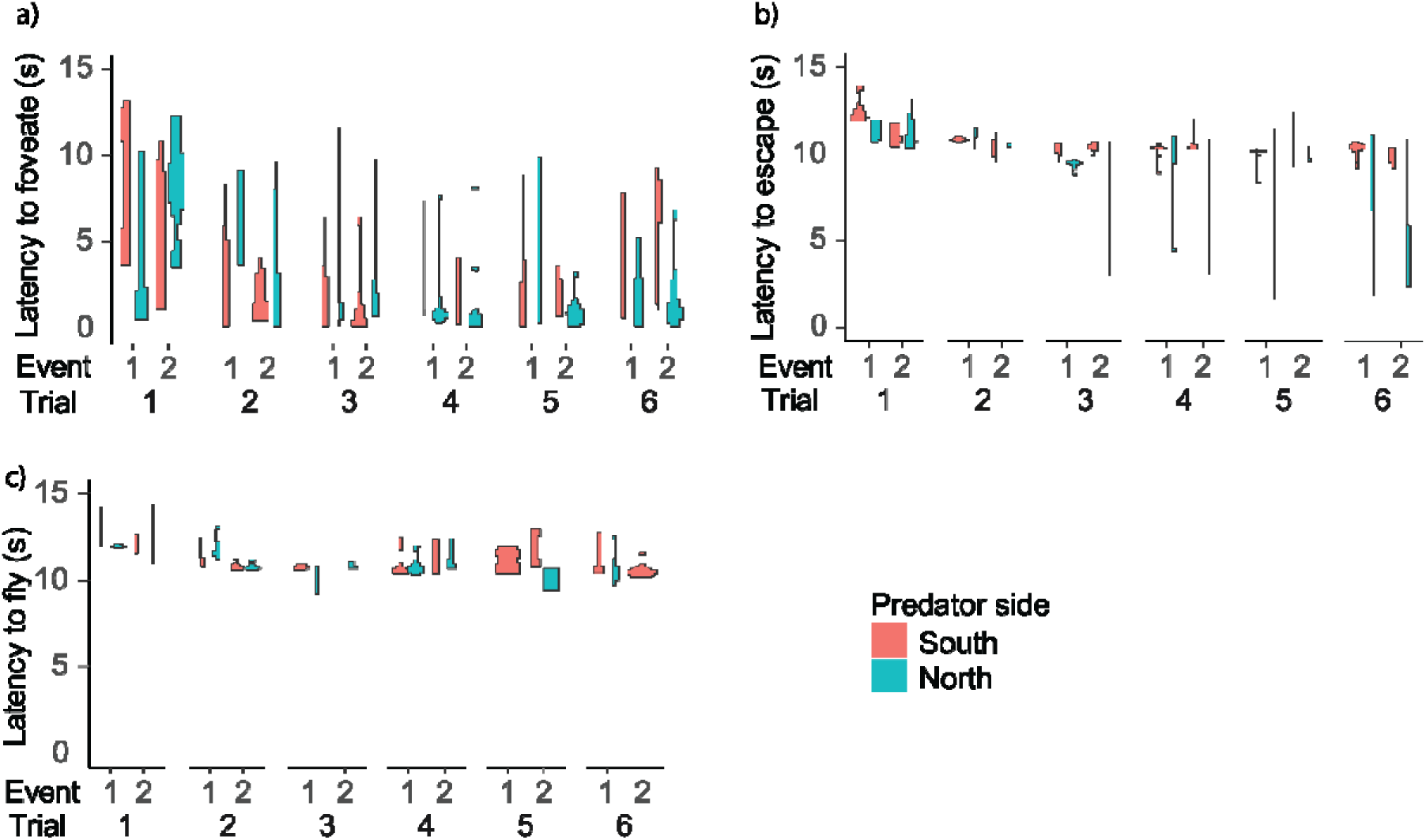
Response variables as a function of the trials and events. a) latency to foveate; b) latency to escape; c) latency to fly. The distribution of the data is displayed as violin plots and the color represents the side of the predator event.

**Figure S2:**
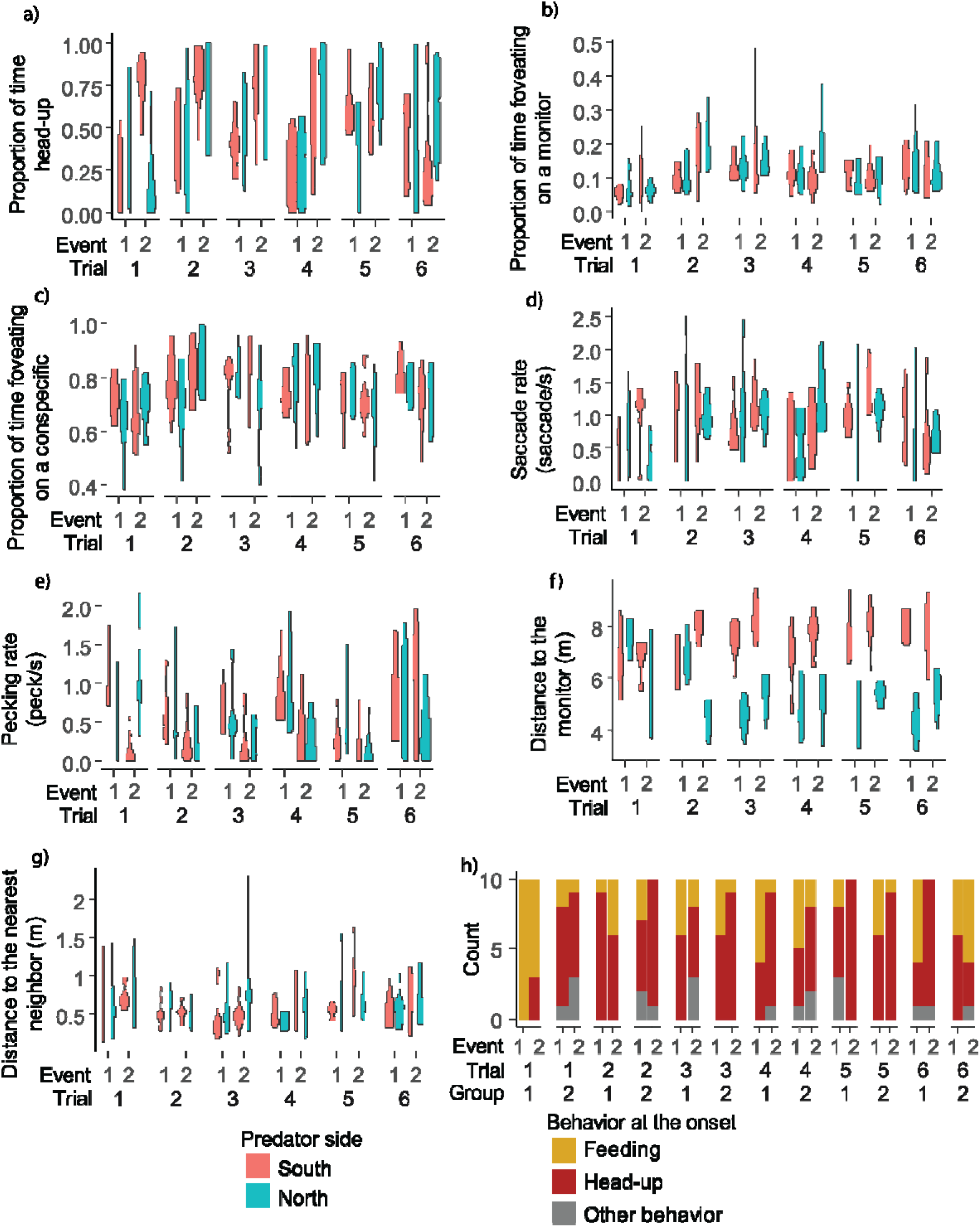
Predictor variables as a function of the trials and events. The distributions of the different variables are displayed as violin plots and the color represents the side of the predator event. a-e) behavior during the “before cue” period (2 minutes period straight before the onset of the looming stimulus) as a function of the trial and event: proportion of time head-up (a), proportion of time foveating on either of the monitors (b), proportion of time foveating on one of the conspecifics (c), saccade rate (d), pecking rate (e). f-h) behavioral and spatial state measures at the onset of the looming stimulus as a function of the trial and event: distance to the monitor (f), distance to the nearest neighbor pigeon (g), the number of individuals within a flock exhibiting respective behaviors (feeding, head-up or other behavior) exhibited at the onset on the looming stimulus (h).

### Definition of object shape

We approximated objects of interest (monitors, hiding places, and other pigeons) as 3D volumes, defining them as spheres or combinations of spheres.

The monitors were represented as a single sphere. The sphere’s center was the monitor’s centroid, and its diameter was set at 86.13 cm, which is the monitor’s diagonal length of 71.77 cm plus a 20% margin. The predator hiding place was also defined as a single sphere, with its center located below the monitor’s centroid at a height of 38 cm above the ground (along the z-axis of the global coordinate system, corresponding to half the height of the hiding place). Its diameter was 158.47 cm, based on the hiding place’s diagonal length of 132.06 cm, plus a 20% margin.

The pigeons were represented by two spheres, denoting the head and the body. The head sphere’s centroid was the midpoint between the eyes, with a diameter of 12 cm (calculated from the distance from the beak tip to the midpoint of the eyes, approximately 4 cm, plus a 2 cm margin for the radius). The body sphere’s centroid was set 6 cm below the center point of the backpack markers (along the Z-axis of the global coordinate system, approximately at the body’s midpoint directly above the pigeon’s paws), and its diameter was 24 cm (considering the centroid was usually around 10 cm above the ground, with an added 20% margin for the radius) (Fig S3).

**Figure S3:**
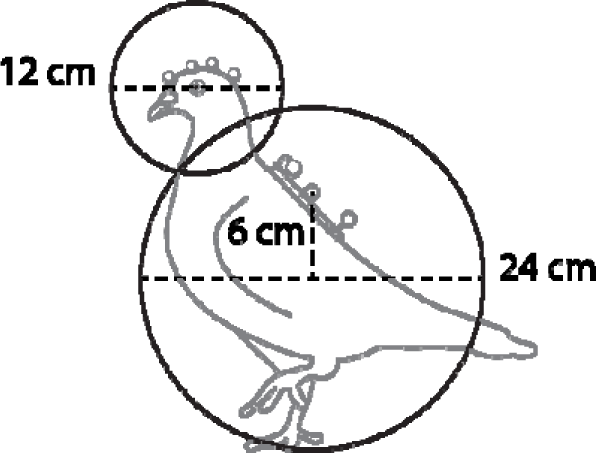
Illustration of the defined spherical representations for the pigeon’s head and body.

### Definition of responses

**Table S2:** Explanation of the different responses and variables used in our models.

| Variable type | Test Set | Variable name | description |
| --- | --- | --- | --- |
| Responses | 1 | Proportion of time foveating | Normalized frequency (normalized to a unit sum) at which each object of interest was observed within the foveal region (in the reprojection of the pigeon's visual field), during the predator event. |
|  | 2.1-2.2 | Latency to foveate (ms) | Latency of the first foveation on the looming stimulus or the predator hiding place after the onset of the looming stimulus and before the predator appears. To exclude the possibility of random gaze crossing, the first foveation was defined as instances lasting longer than 300 ms, which occurred at least 200 ms after the onset of the looming stimulus. |
|  | 3-4.1 | Escaped before the looming stimulus offset or not (binary) | Whether the first overt escape response (either running away or flying) occurred before the looming stimulus offset (1) or not (0; i.e., either the pigeon responded after the looming stimulus offset or did not exhibit a detectable response). |
|  | 3-4.1 | Latency to fly (ms) | Latency of the first flight after the onset of the looming stimulus |
| Predictors | 1 | Object type | Type of object: the monitor displaying the looming stimulus (relevant environmental cue), the monitor not displaying the looming stimulus (irrelevant environmental cue), or the nearest neighbor (conspecific) |
|  | 2.1 | Behavior at the onset | Behavior at the onset of the looming stimulus (either head-up or feeding, other behaviors are not considered as they occurred much less often (20 occurrences out of 240 observations) |
|  | 2.1 | Distance from the monitor (mm) | Distance to the monitor at the onset of the looming stimulus |
|  | 2.1 | Distance to nearest neighbor (mm) | Distance to the nearest neighbor at the onset of the looming stimulus |
|  | 2.2 | Head-up | Proportion of time the pigeon spent being head-up during the 2-min “before-cue” period |
|  | 2.2 | Foveate on the monitor | Proportion time spent foveating on either monitor during the 2-min “before-cue” period |
|  | 2.2 | Foveate on any pigeon | Proportion time spent foveating on any pigeon during the 2-min “before-cue” period |
|  | 2.2 | Saccade rate (saccades/s) | Saccade rate when the pigeon is head-up during the 2-min “before-cue” period |
|  | 2.2 | Pecking rate (pecks/s) | Pecking rate during the 2-min “before-cue” period |
|  | 3 | Latency to foveate (ms) | See above |
|  | 4.1 | First latency to foveate (ms) | The latency to foveate on the predator cue by the first pigeon during the looming stimulus presentation |

### Automated behavior classification and rating

Behaviors were automatically classified using postural definitions on the 3D data from the motion capture system (Table S3).

**Table S3:** Description of the behavior classification definitions. Terms definitions: state = behavior that lasts in time; event = punctual behavior. Body-head vector = the vector originating from the body center and projecting to the head centroid. Head direction vector = the vector projecting to the front of the head and towards the horizon when the head is still (corresponds to the y axis of the local coordinate system, projecting 30° above the beak-head center axis). Pitch = head rotation movement corresponding to “nodding” (pitch = 0° corresponds to the head direction vector pointing towards horizon, pitch < 0° corresponds to the head direction vector pointing down, pitch > 0° corresponds to the head direction vector pointing up). Roll = head rotation movement where the head is “tilted” (roll = 0° corresponds to a straight head and roll ≠ 0° corresponds to a head tilted to the right or to the left).

| Behavior | Category | Definition |
| --- | --- | --- |
| head-down | state | The centroid of the head (middle point between the eyes) is lower than the centroid of the body (6cm below the backpack markers center point) along the z axis of the global coordinate system (=vertically) |
| head-up | state | The pigeon is not head-down and not exhibiting another behavior (feeding, grooming or courting) |
| pecking | event | The beak tip is closer than 4cm from the ground and is not oriented towards the body (pitch > -100° from horizon). The pecking event is defined as the frame where the beak tip is the closest from the ground |
| feeding | state | Any time between 2 pecks spaced no more than 6 seconds apart |
| grooming | state | Preening the back feathers: This occurs when the head is pointing backward (defined as when the head direction vector and the body-head vector form an angle larger than 60°) and the beak tip is close to the body center (less than 10cm) OR when the head is pointing backward (defined as when the head direction vector and body-head vector form an angle larger than 40°) and the beak tip is close to the body centroid (less than 10cm), and the head centroid is above the body centroid.<br>Preening the breast feathers: This behavior occurs when the head is strongly pointing down (pitch < -100°) OR when the head is pointing down (pitch < -80°) and the head direction vector and the body-head vector form an angle larger than 60°.<br>Scratching the head with the paw: This behavior is exhibited when the roll of the head is large (> 50°) and the head is low (less than 3cm above the body center). |
| courting | state | The pigeon is bowing its head, which is a typical head movement in pigeon courtship, at a rate of at least 2 bows within an 8-second period, while being within a proximity of less than 60cm and oriented towards another pigeon within an angle of less than 120°. We filtered out any instances shorter than 1 second for robust detection. |
| running away from the monitor | state | When the body centroid of the pigeon moves at a speed faster than 0.6m/s, and the pigeon is not classified as courting or being courted, while also increasing its distance from the monitor at a speed of 0.2m/s, we filter out any events shorter than 70ms for robust detection. |
| flying | state | When the body centroid of the pigeon moves at a speed greater than 2m/s, we filter out any events shorter than 70ms for robust detection. |

The reliability of the automated classification was verified by two human raters through manual coding. This process was conducted on a dataset different from the one used in this study. The raters assessed the presence or absence of specific behaviors (or the number of pecks) in 1-second sample segments (for coding peck counts, 3-second segments were used to increase variability in the number of pecks). We coded 50 segments for each behavior, with the exception of flying (for which only 30 segments were coded due to its scarcity in the dataset) and pecking (for which 60 segments were coded to enhance variation in peck counts). Behaviors such as head-down, head-up, and feeding were not rated, as they were determined based on a simple threshold definition and/or depended on rated behaviors. Inter-rater reliability was calculated using R (R Core Team, 2022) and the ‘irr’ package (Gamer et al., 2019). The pecking count IRR was computed using Intraclass Correlation Coefficient (ICC) for count data, while grooming, courting, and flying were analyzed using Cohen’s kappa for binary, presence-absence data.

The head-up/down state was defined based on the relative position of the head and body. The head-up state was further refined to exclude periods of feeding, grooming, and courting, to avoid moments when pigeons were likely distracted by these activities. To verify the detection of pecks, two human raters counted the number of pecks in short data segments. The Intraclass Correlation Coefficient (ICC) confirmed a very high agreement between their coding and automated detection (ICC_min = 0.98; see Table S4). Feeding was then defined based on consecutive pecking instances. Grooming and courting were identified by a combination of several postural states and validated by the same two human raters (presence or absence of the behavior in short data segments; Cohen_’_s kappa_min = 0.8; see Table S4). Running was defined based on a speed threshold, determined by inspecting locomotive speed histograms (Fig S4). This definition was limited to pigeons moving away from the predator hiding place and not engaged in courting activities, to identify ‘running away from the monitor’ as an evasion response. Flying was observable when a pigeon took off but was specifically defined by the locomotive speed exceeding 2 m/s, a threshold unattainable by running alone. This threshold was further validated by the human raters’ coding (presence or absence of the behavior in short segments; Cohen’s kappa_min = 0.93; see Table S4). All rated behaviors achieved Cohen’s Kappa or Intraclass Coefficient (ICC) scores above 0.8 (“almost perfect”; (Landis & Koch, 1977), indicating a strong agreement between human raters and the automated behavior coding.

**Table S4:** Inter-rater reliability for behavior classification. Pecking count was rated with ICC and other behaviors with Cohen’s kappa. HR1 and HR2 represent the human raters and ACC the automated computer classification.

| Behavior | Inter-rater reliability |  |  | Sample number | Metric | Comparison method |
| --- | --- | --- | --- | --- | --- | --- |
|  | HR1-ACC | HR2-ACC | HR1-HR2 |  |  |  |
| Pecking | 0.983 | 0.984 | 0.987 | 60 | count | ICC |
| Grooming | 1 | 0.96 | 0.96 | 50 | presence-absence | Cohen's Kappa |
| Courting | 0.88 | 0.8 | 0.838 | 50 | presence-absence | Cohen's Kappa |
| Flying | 0.933 | 1 | 0.933 | 30 | presence-absence | Cohen's Kappa |

**Figure S4:**
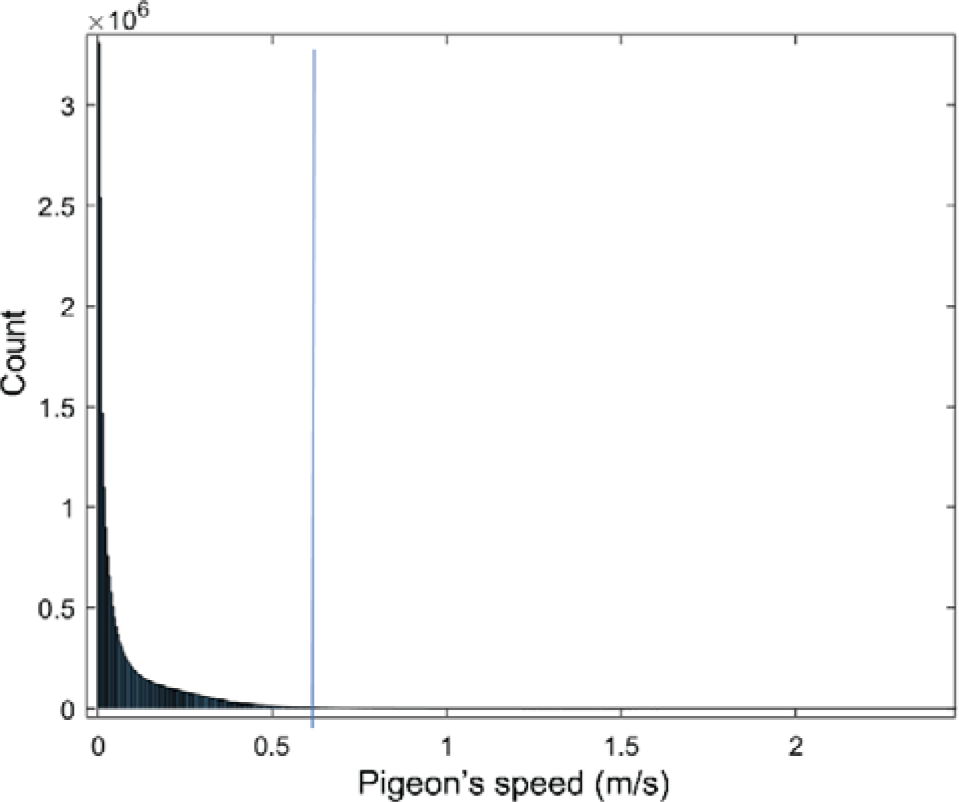
Histogram of the pigeons’ speed for the whole dataset. Running unusually fast is defined as the body center moving with a speed larger than 0.6 (vertical blue line)

### Behavioral changes after the predator presentation event

We investigated whether various behavioral and spatial measures were influenced by a predation event. We collected data for all measures over a one-minute period immediately preceding the onset of the looming stimulus and following the predator’s disappearance (averaging the distances over the same period). Linear Mixed Models (LMMs) were used, with the behavior/distance as the response variable. The time period (before vs. after the event) served as the test variable, while event type, predator side, and sex were control variables. Pigeon ID and trial were included as random effects. For all tested behaviors, measurements taken after the predation event showed significant differences compared to those before (p < 0.001). The distances also exhibited significant changes (distance from the closest monitor: p = 0.0034; distance from the nearest neighbor: p = 0.0043; see Fig S5 for a graphical representation and Table S5 for detailed model results).

**Figure S5:**
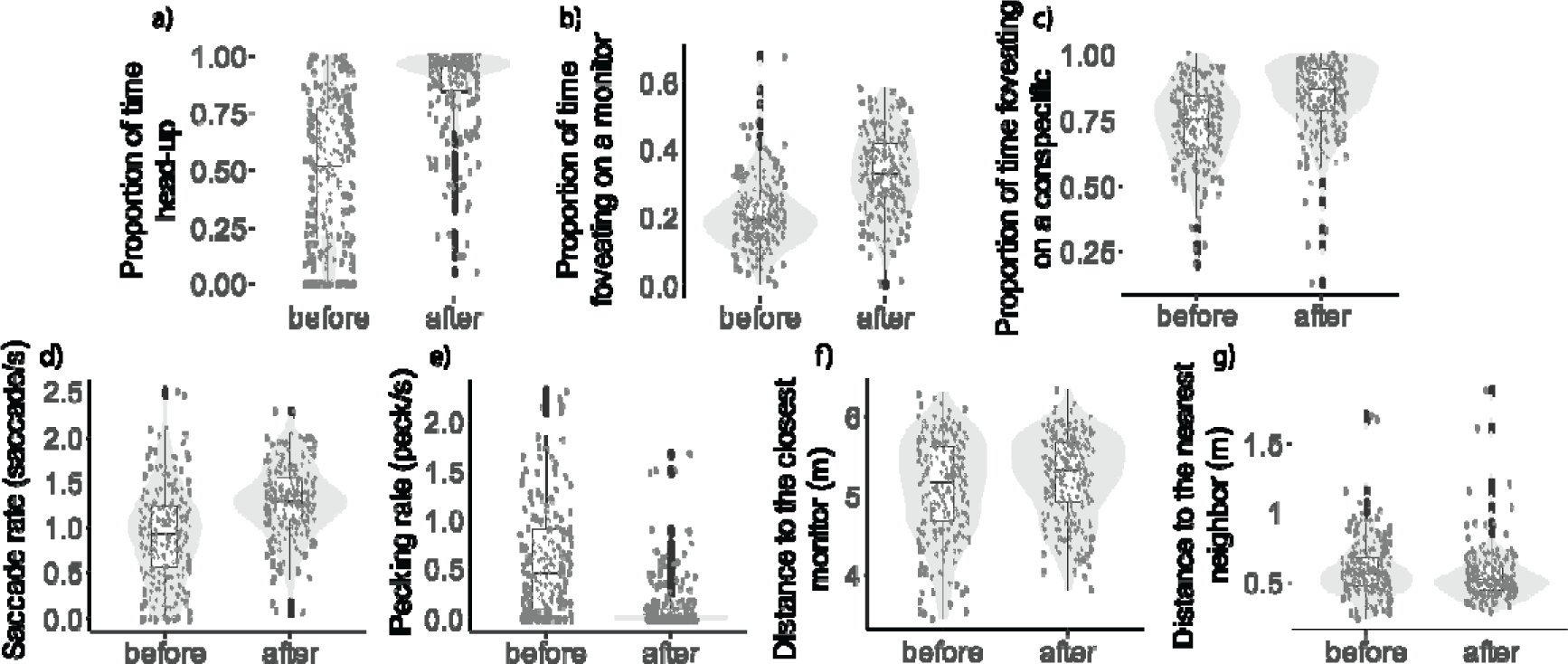
Behavioral changes between the 1 min period before the onset of the looming stimulus and the 1 min period after the disappearance of the predator: proportion of time spent being head-up (a), proportion of time spent foveating on one of the monitors (b), proportion of time spent foveating on one of the conspecifics (c), saccade rate (d), pecking rate (e), distance to the closest monitor (f), distance to the nearest neighbor (g). The dots represent single observations (jittered horizontally for visualization), the grey shade represents the violin plot of the distributions, and the boxplot’s boxes represent the 0.25 and 0.75 quartiles (with the median represented as a line inside the box) and the whiskers the minimum and maximum values within the lower/upper quartile ± 1.5 times the interquartile range.

**Table S5:** Model results for behavioral comparison between the 1 min period before the onset of the looming stimulus and the 1 min period after the disappearance of the predator. For all models, the table details the beta estimates, as well as the Chi square value, degrees of freedom and p-values from the likelihood ratio test. Event, predator side, sex, pigeon ID (random effect) and trial (random effect) has been added as control variables.

| Response | Type | Variable | # of obs. | Beta | Chi-sq | Df | p |
| --- | --- | --- | --- | --- | --- | --- | --- |
| Head-up | Test variable | Time period | 477 | 0.27 | 237.71 | 1 | <0.00001 |
|  | Control variable | Event | 477 | 0.13 | 33.39 | 1 | <0.00001 |
|  |  | Predator side (North) | 477 | 0.00 | 0.05 | 1 | 0.832 |
|  |  | Sex (male) | 477 | -0.08 | 3.18 | 1 | 0.07433 |
| Pecking rate (log-transformed) | Test variable | Time period | 477 | 0.27 | 237.71 | 1 | <0.00001 |
|  | Control variable | Event | 477 | 0.13 | 33.39 | 1 | <0.00001 |
|  |  | Predator side (North) | 477 | 0.00 | 0.05 | 1 | 0.832 |
|  |  | Sex (male) | 477 | -0.08 | 3.18 | 1 | 0.07433 |
| Saccade rate | Test variable | Time period | 477 | 0.27 | 94.75 | 1 | <0.00001 |
|  | Control variable | Event | 477 | 0.09 | 5.49 | 1 | 0.01907 |
|  |  | Predator side (North) | 477 | -0.02 | 0.35 | 1 | 0.55565 |
|  |  | Sex (male) | 477 | -0.03 | 0.15 | 1 | 0.69753 |
| Proportion of time looking at any monitor | Test variable | Time period | 477 | 0.09 | 174.32 | 1 | <0.00001 |
|  | Control variable | Event | 477 | 0.02 | 6.09 | 1 | 0.01357 |
|  |  | Predator side (North) | 477 | 0.00 | 0.01 | 1 | 0.93284 |
|  |  | Sex (male) | 477 | -0.04 | 2.47 | 1 | 0.11573 |
| Proportion of time looking at any conspecific | Test variable | Time period | 477 | 0.07 | 76.63 | 1 | <0.00001 |
|  | Control variable | Event | 477 | 0.00 | 0.09 | 1 | 0.77029 |
|  |  | Predator side (North) | 477 | -0.01 | 0.89 | 1 | 0.34477 |
|  |  | Sex (male) | 477 | 0.04 | 2.48 | 1 | 0.11564 |
| Distance to the closest monitor | Test variable | Time period | 477 | 101.61 | 8.58 | 1 | 0.00339 |
|  | Control variable | Event | 477 | 9.47 | 0.04 | 1 | 0.84643 |
|  |  | Predator side (North) | 477 | 51.44 | 1.11 | 1 | 0.2922 |
|  |  | Sex (male) | 477 | 8.38 | 0.01 | 1 | 0.91392 |
| Distance to the nearest neighbor | Test variable | Time period | 477 | -31.11 | 8.15 | 1 | 0.0043 |
|  | Control variable | Event | 477 | 6.07 | 0.16 | 1 | 0.69182 |
|  |  | Predator side (North) | 477 | 26.76 | 3.03 | 1 | 0.08195 |
|  |  | Sex (male) | 477 | -50.13 | 1.84 | 1 | 0.17515 |

### R formula

**Table S6:**
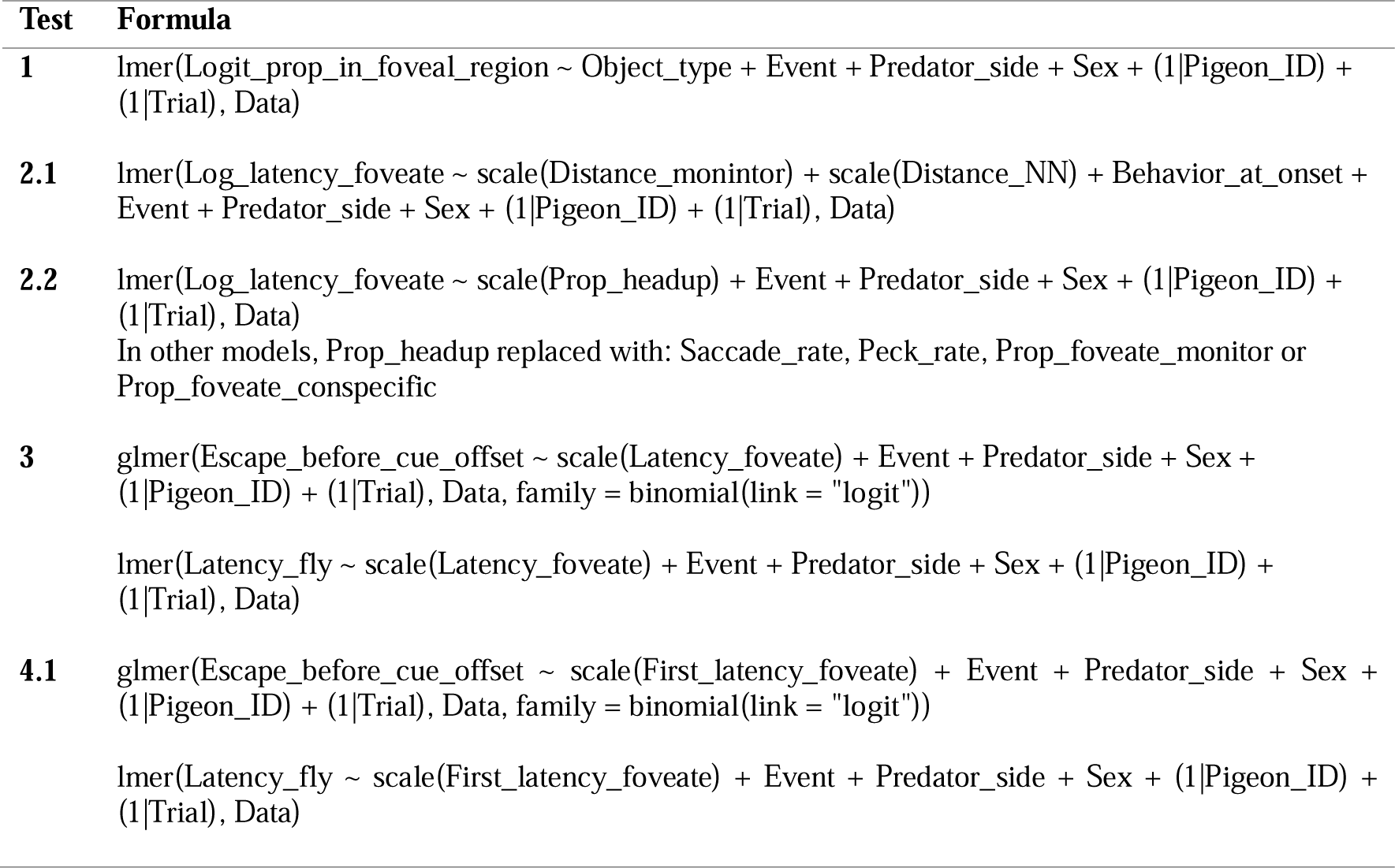
R formulas used for the different models. Variables starting with “Logit” have been logit-transformed, with “Log” have been log-transformed. As control variables, we have included in all models the event number, the predator side, the sex of the individual, the pigeon ID (random effect) and the trial ID (random effect).

### Results of main models

**Table S7:**
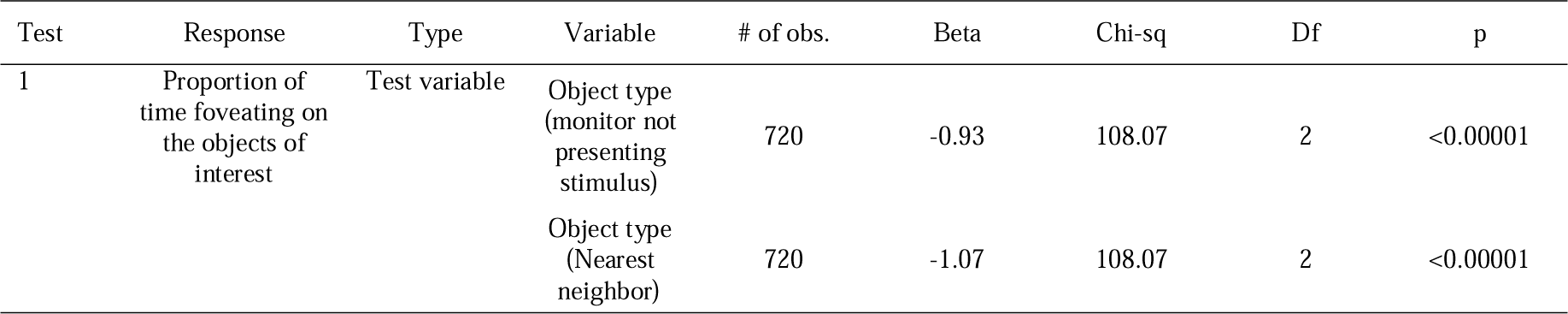
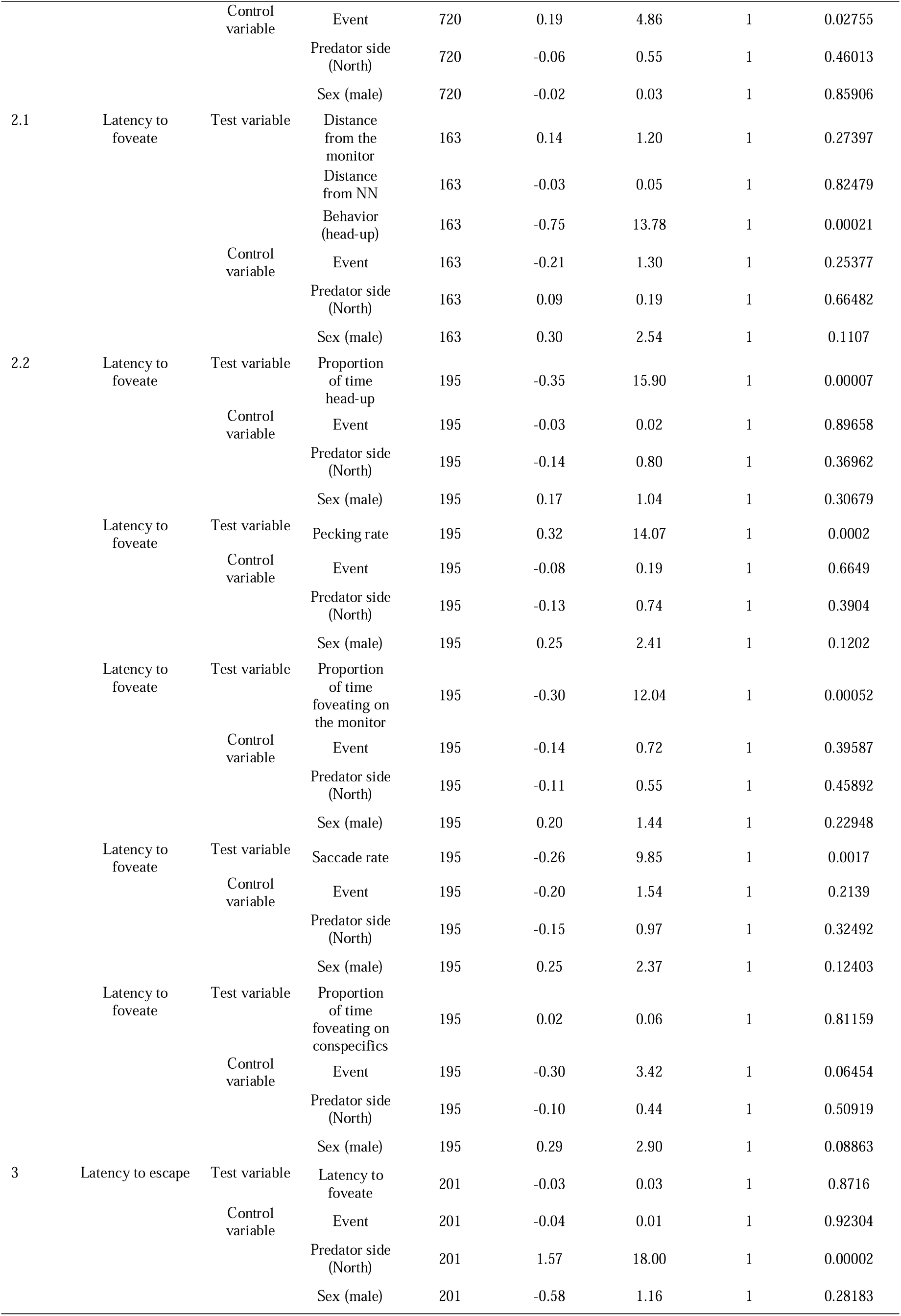
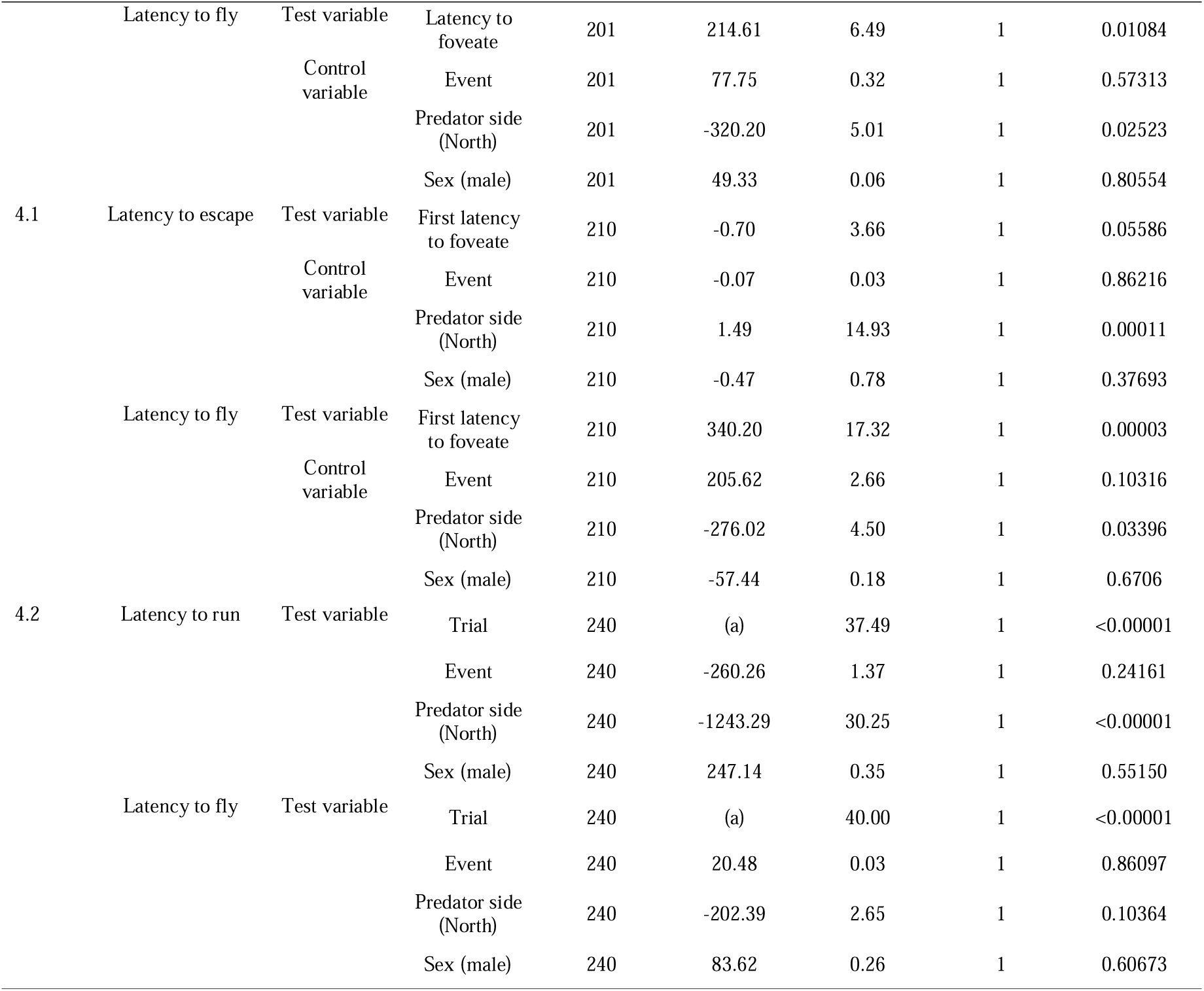
Results of all models related to our main hypotheses. This table presents detailed information for each predictor in every model used in this study, including the beta estimates from the model, as well as the Chi-square value, degrees of freedom, and p-values derived from the likelihood ratio test. Note: (a) was incorporated as a random effect in the model and, as such, does not have a beta estimate.

### Repeatability analysis

We assessed whether the variables used in the models (both responses and predictors) exhibited significant inter-individual differences by evaluating their repeatability across trials and events. The rpt() function from the rptR package in R (Stoffel et al., 2017) was utilized for this analysis, incorporating the event, predator side, and the sex of the individual as control variables, and the trial as an additional random effect in the repeatability model. Of the three response latencies examined, only the latency to escape demonstrated significant repeatability (R = 0.15, p < 0.0001), suggesting that certain individuals consistently escaped faster than others. In examining the response delay, we specifically analyzed the response time delay for flights (latency to fly -latency to foveate), which was found to be significantly repeatable. However, neither the latency to foveate nor the latency to fly were repeatable on their own. Due to the observation that in many trials, some pigeons did not fly up during the predator presentation, we also investigated whether the decision to fly was repeatable. Our analysis revealed that certain pigeons were more prone to flying than others (R = 0.22, p < 0.0001). As for the predictors tested, nearly all were found to be significantly repeatable (R ranging from 0.06 to 0.17, p values all below 0.0362), except for the distance from the monitor (p = 1). Detailed results of this analysis are presented in Table S8.

**Table S8:** Results of all repeatability analyses including the observed variable, the repeatability estimate and the p-value.

| Variable | Repeatability estimate | p-value |
| --- | --- | --- |
| Latency to foveate (log-transformed) | 0.011 | 0.359 |
| Latency to escape | 0.145 | <0.00001 |
| Latency to fly | 0.0525 | 0.062 |
| Response time delay between foveation and flight | 0.128 | 0.0404 |
| Flew or not | 0.219 | <0.00001 |
| Proportion of time head-up | 0.166 | <0.00001 |
| Saccade rate | 0.126 | 0.00014 |
| Pecking rate (log-transformed) | 0.0919 | 0.00507 |
| Proportion foveate on the predator cue | 0.144 | 0.00001 |
| Proportion foveate on conspecifics | 0.132 | 0.00007 |
| Distance from the monitor | <0.0001 | 1 |
| Distance from nearest neighbor | 0.0644 | 0.0362 |

### Supplementary graphs of Test 2.2

**Figure S6:**
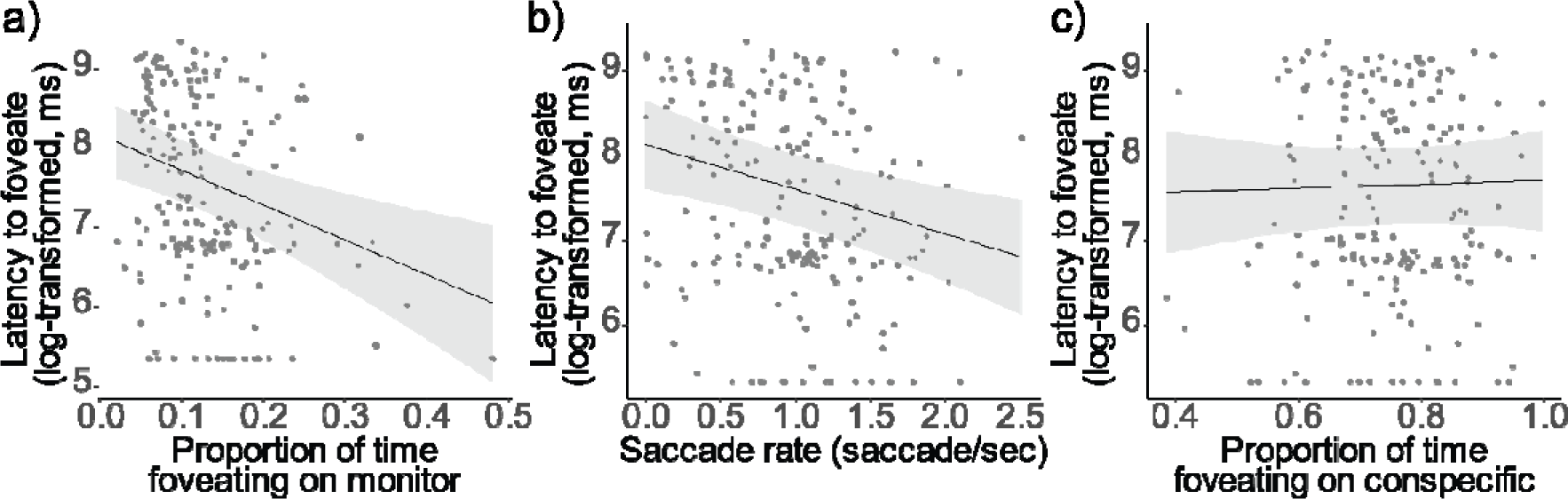
Results of the models from Test 2.2: Latency to foveate on the predator cue as a function of the proportion of time foveating on the monitor during the “before-cue” period (a), as a function of the saccade rate during the same period (b) and the proportion of time foveating on any conspecific during the same period (c). The regression lines were determined with other variables held constant, set to their mean values. A comprehensive table detailing the outcomes of the models can be found in Table S7.

